# HMGB1 coordinates SASP-related chromatin folding and RNA homeostasis on the path to senescence

**DOI:** 10.1101/540146

**Authors:** Konstantinos Sofiadis, Milos Nikolic, Yulia Kargapolova, Natasa Josipovic, Anne Zirkel, Antonis Papadakis, Ioanna Papadionysiou, Gary Loughran, James Keanes, Audrey Michel, Eduardo G. Gusmao, Athanasia Mizi, Theodore Georgomanolis, Janine Altmüller, Peter Nürnberg, Andreas Beyer, Argyris Papantonis

## Abstract

Spatial organization and gene expression of mammalian chromosomes are maintained and regulated in conjunction with cell cycle progression. This is however disturbed once cells enter senescence and the highly abundant HMGB1 protein is depleted from senescent cell nuclei to act as an extracellular proinflammatory stimulus. Despite its physiological importance, we know little about the positioning of HMGB1 on chromatin or about its roles in the nucleus. To address this, we mapped HMGB1 binding genome-wide in different primary cells using a tailored protocol. We integrated ChIP-seq and Hi-C data with a graph theory approach to uncover HMGB1 demarcation of a subset of topologically-associating domains (TADs) that harbor genes required for paracrine senescence. Moreover, using sCLIP, knock-down and overexpression experiments, we now show that HMGB1 is a *bona fide* RNA-binding protein (RBP) bound to senescence-relevant mRNAs and affecting splicing. HMGB1 also has an interactome rich in RBPs, many of which are implicated in senescence regulation. The mRNAs of many of these RBPs are directly bound by HMGB1 and concertedly regulate the availability of SASP-relevant transcripts. Our findings highlight a broader than hitherto assumed role for HMGB1. It coordinates chromatin folding and RNA homeostasis as part of a feedforward loop controlling both cell-autonomous and paracrine senescence inside and outside of cells.

## Introduction

The high-mobility group box 1 (HMGB1) protein, a member of the highly conserved non-histone DNA binding HMG protein family, was named after its characteristically rapid electrophoretic mobility (Štros, 2010). HMGB1 is the most abundant non-histone protein in mammalian nuclei, with 1 HMGB1 molecule per every ~10 nucleosomes (Thomas and Stott, 2012). Despite its high abundance and conservation, HMGB1 has been predominantly studied as an extracellular signaling factor, hence its characterization as an “alarmin” (Lohani and Rajeswari, 2016; Bianchi *et al*, 2017).

To function as an alarmin, HMGB1 is actively secreted by cells like activated monocytes and macrophages or passively released by necrotic and damaged cells. Once received by other cells in the niche, HMGB1 is recognized by RAGE receptors to potently signal inflammation (Scaffidi *et al*, 2002; Bonaldi *et al*, 2003; Orlova *et al*, 2007). In cells entering senescence, HMGB1 translocates from the nucleus to the cytoplasm and is then secreted to stimulate NF-κB activity via Toll-like receptor signaling. This relocalization and secretion controls the senescence-associated secretory phenotype (SASP) of mammalian cells, thus representing a major paracrine contributor both *in vitro* and *in vivo* (Salminen *et al*, 2012; Acosta *et al*, 2013; Davalos *et al*, 2013).

Inside proliferating cell nuclei, HMGB1 has been studied in some detail for its contribution to DNA repair (Ito *et al*, 2015; Mukherjee *et al*, 2016), V(D)J recombination (Little *et al*, 2013; Zagelbaum *et al*, 2016) or chromatin assembly (Bonaldi *et al*, 2002), but far less for its transcriptional role (Calogero *et al*, 1999; Mitsouras *et al*, 2002). Cells lacking HMGB1 contain reduced numbers of nucleosomes, rendering chromatin more susceptible to DNA damage, spurious transcription, and inflammatory activation (Giavara *et al*, 2005; El Gazzar *et al*, 2009; Celona *et al*, 2011; De Toma *et al*, 2014). As regards its association with chromatin, HMGB1 is thought to bind in a nonspecific manner via its two HMGB-box domains. This allows it to bend and contort DNA and, thus, to facilitate recruitment of transcription factors like p53 (Štros, 2010; Rowell *et al*, 2012). HMGB1 associates with cognate DNA sites via characteristically high “on/off” rates, and its acidic tail is important for stabilizing binding (Pallier *et al*, 2003; Ueda *et al*, 2004; Štros, 2010; Blair *et al*, 2016). However, HMG-box DNA-binding domains are particularly insensitive to standard fixatives like formaldehyde (Pallier *et al*, 2003; Teves *et al*, 2016). Thus, capturing HMGBs on chromatin is challenging, and there exist no genome-wide datasets describing HMGB1 binding in mammalian cells (see http://chip-atlas.org). As a result, our appreciation of its on-chromatin roles remains vague.

To address this, we employ here a tailored approach that previously allowed us to efficiently map HMGB2 binding sites genome-wide (Zirkel *et al*, 2018). We can now show that HMGB1 binding in primary endothelial and lung fibroblast cells is far from nonspecific, while also disparate to that by HMGB2. Following integration of its binding positions with genome-wide chromosome conformation capture data (Hi-C; Kempfer and Pombo, 2019), we found that HMGB1 demarcates the boundaries of a considerable and specific fraction of topologically-associating domains (TADs; Dixon *et al*, 2012; Nora *et al*, 2017). This topological contribution is eliminated upon senescence entry, and knockdown/ overexpression experiments show that HMGB1 controls the expression of genes that are central to the SASP and are specifically embedded in these TADs. Critically, as HMGB1 was proposed to have RNA-binding capacity (Castello *et al*, 2016), we used sCLIP (Kargapolova *et al*, 2017) to show it also influences the splicing and stability of senescence-relevant mRNAs. This occurs via a network of RNA-binding factors that interact with HMGB1 and are also under its direct control. In summary, using replicative senescence as a model, we characterize the multifaceted *in nucleo* roles of HMGB1 that converge on the coordination of chromatin and RNA control for SASP regulation.

## Results

### Senescence entry is marked by HMGB1 nuclear loss and cell secretion

To investigate the nuclear roles of HMGB1 across cellular contexts, we used primary human umbilical vein endothelial cells (HUVECs) and fetal lung fibroblasts (IMR90) that are of distinct developmental origins and have disparate gene expression programs. We defined an early-passage proliferative state and a late-passage senescent state by combining β-galactosidase staining, cell cycle staging by FACS, and MTT proliferation assays (**Fig. 1A**), as well as a “senescence clock” based on the methylation of six CpGs (Zirkel *et al*, 2018). Next, we used RNA-seq data from proliferating and senescent HUVEC and IMR90 (from two different donors/isolates) to look into changing mRNA levels of chromatin factors. Significant and convergent changes between the two cell types involved strong suppression of histone chaperones and histones, lamin-associated proteins, centromere components, cohesin and condensin complexes, as well as all of HMGB/N-family proteins (**Fig. 1B**). Many of these factors were consistently suppressed also at the protein level (**Fig. 1C**; Davalos *et al*, 2013; Shah *et al*, 2013; Rai *et al*, 2014; Zirkel *et al*, 2018). We chose to focus on HMGB1 due to its high conservation, nuclear abundance (Thomas and Stott, 2012; **Fig. 1D**) and key role in SASP induction (Davalos *et al*, 2013), but mostly due to its still elusive function on chromatin, especially in respect to spatial chromosome organization.

**Figure 1.**
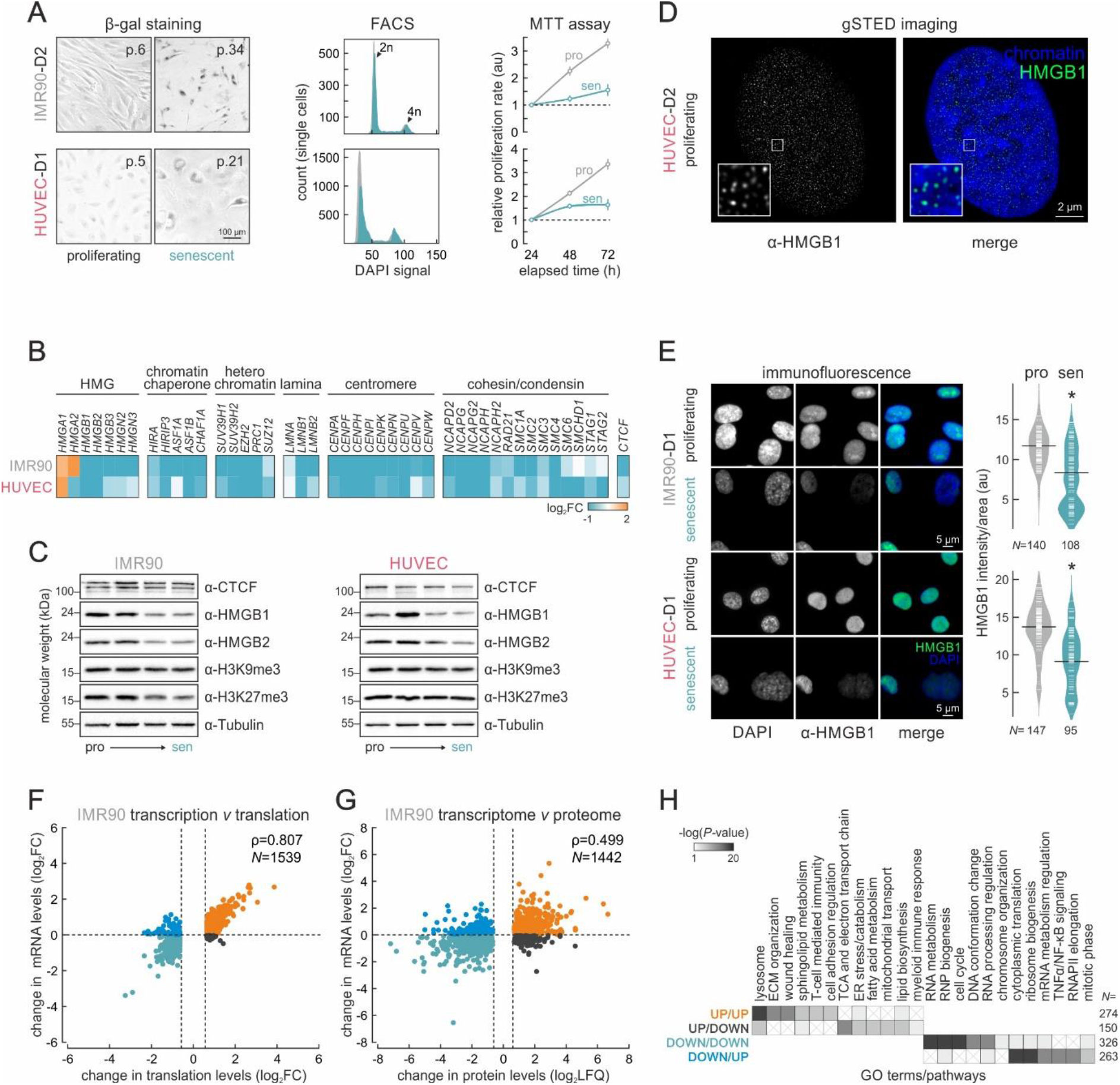
Senescence entry is transcriptionally driven in primary human cells. (A) Proliferating and senescent IMR90 and HUVECs assayed for β-galactosidase activity (*left*), cell cycle profiling via FACS (*middle*) and proliferation via MTT assays (*right).* (B) Heatmaps showing changes in gene expression levels upon senescence (log_2_FC) of genes encoding selected chromatin-associated factors. For each gene shown, statistically significant expression changes were recorded in at least one cell type. (C) Western blots showing changing protein levels on the path to senescence in IMR90 and HUVECs. Passage 6 cells represent the proliferating state, and passages 21 and 32 the senescent state for HUVECs and IMR90, respectively. (D) Super-resolution (gSTED) imaging of HMGB1 distribution in proliferating HUVEC nuclei counterstained with DAPI. Bar: 2μm. (E) Representative immunofluorescence images of IMR90 and HUVECs (*left*) show reduced HMGB1 levels in senescent nuclei; bean plots quantify this reduction (*right; N* indicates the number of cells analyzed per each condition/cell type). Bars: 5 μm. *:*P*<0.01; Wilcoxon-Mann-Whitney test. (F) Scatter plots showing correlation between RNA-seq (transcription) and Ribo-seq (translation; log_2_) in proliferating and senescent IMR90. Pearson’s correlation values (ρ) and the number of genes in each plot (*N*) are also shown. (G) As in panel F, but for the correlation between RNA-seq and whole-cell proteome changes. (H) Heatmap showing GO terms/pathways associated with gene subgroups from panel G. The number of genes in each subgroup (*N*) are also shown.

Immunodetection in early- and late-passage cells documented a >50% decrease in HMGB1 nuclear levels in the heterogeneous senescence-entry populations of HUVEC or IMR90 (**Fig. 1E**). HMGB1 nuclear depletion was most dramatic in the enlarged nuclei of senescent cells of either cell type, while smaller nuclei remained largely unaffected. FACS-sorting IMR90 based on light scattering allowed enrichment for cell populations with enlarged nuclei (i.e., ~70% of cells had larger than average nuclei, with >35% being >1.5-fold larger than the average proliferating nucleus; **Fig. S1A**). This showed that enlarged nuclei lacking HMGB1 almost invariably represent senescent cells and exhibit a drop in the H3K27me3 mark of facultative heterochromatin – effects which would otherwise be masked (**Figs 1C** and **S1B,C**). Last, we showed that it is those larger cells that secrete HMGB1, but not HMGB2, into their growth medium presumably to contribute to paracrine senescence via the SASP (**Fig. S1D**).

### The senescence program is transcriptionally-driven down to the single-cell level

Despite strong changes documented by RNA-seq, it is still not known to which extent the senescent program is implemented via changes at the transcriptional or the translational level. To address this, we generated matching mRNA-seq, Ribo-seq and whole-cell proteomics data from proliferating and senescent IMR90 in biological triplicates. Comparative analysis of mRNA-seq and Ribo-seq data showed that essentially all significant changes at the level of mRNA translation were underlied by equivalent changes in transcript availability (**Fig. 1F**). Only few transcripts (~800) showed increased translation to counteract transcriptional suppression (e.g., *TNFRSF19, HMGN2, LMNB2)* or the converse (e.g., *IL12A, CDKN1A, CDKN2B).* Gene set enrichment analysis of the two subgroups (with a “buffer” of at least log_2_ 0.6) showed that transcripts translationally upregulated while transcriptionally suppressed are linked to the formation and secretion of endosomal vesicles (and thus to HMGB1 release into the extracellular milieu). On the other hand, transcripts downregulated as regards translation yet transcriptionally upregulated associate with ribosomal complex formation and translation, but surprisingly also with RNA binding and RNA catabolism (**Fig. S1E**).

Similar analysis of mRNA-seq against whole-cell proteome data also showed that the vast majority of proteins with significantly altered levels in senescence were also equally regulated at the level of transcription (**Fig. 1G**). Genes linked to the major GO terms and pathways characterizing senescence entry were convergently up- (e.g., ECM organization, lysosome) or downregulated at both the mRNA and protein levels (e.g., cell cycle, DNA conformation change, RNA processing; **Fig. 1H**). Curiously, several processes relevant to the SASP (e.g., NF-κB signaling, mRNA metabolism appeared to be regulated by combining higher transcription with diminished protein availability (**Fig. 1H**).

Although predominantly under transcriptional control, senescence entry is idiosyncratic to individual cells and occurs asynchronously in a given cell population. Thus, it is important to obtain a single-cell understanding of the transcriptional changes linked to this transition that is marked by the nuclear loss of HMGB1. To this end, we developed a new protocol for single-cell sequencing of nascent RNA. Typically, scRNA-seq relies on capturing and barcoding cellular mRNAs via reverse-transcription of their poly(A) tails (See *et al*, 2018). To capture nascent RNA instead, we sought to add poly(A) tails to RNAs associated with the active sites of transcription *in situ.* We used our established “factory RNA-seq” protocol (Caudron-Herger *et al*, 2015) as a basis for isolating intact nuclei rich in nascent transcripts from both proliferating and senescent HUVEC. Most chromatin not associated with active transcription sites was removed by DNase digestion and nascent transcripts were polyadenylated *in situ*, before standard library preparation on a 10X Genomics platform (**Fig. S2A**). Using this new approach, and despite lower rates of non-duplet capturing (see **Methods** for details), we analyzed >600 single nuclei carrying an average of 1,650 transcripts each. More than 55% of these transcripts represented introns or intron-exon junctions. Unsupervised t-SNE clustering of these single-nucleus nascent transcriptomes grouped senescent cells in a single cluster, while subdividing the proliferating HUVEC population into three partially overlapping clusters (**Fig. S2B**). Reassuringly, genes differentially-expressed amongst the proliferating and senescent clusters associated with GO terms relevant to senescence and the SASP (**Fig. S2C**). Mapping nascent RNA levels of individual genes onto those clusters, showed that *HMGB1* and *HMGB2* are actively repressed not only in senescent cells, but also in numerous proliferating cells (in line with our previous observations on mRNA levels; Zirkel *et al*, 2018). Conversely, the senescence-induced *HMGA1/A2* loci strongly produce nascent RNA in senescent cells, but are also upregulated in those proliferating cells that show diminished *HMGB1/B2* transcription (e.g., cells of the *blue* cluster; **Fig. S2B,D**). SASP-related genes like *IL4R* and *MMP14* also show strong transcription in cells repressing *HMGB1/B2*, while the p21-encoding locus, *CDKN1A*, has most nascent RNA detected in the senescent cluster (**Fig. S2D**). This implied that the gradual loss of *HMGB* expression marks a temporally-regulated path towards senescence for each cell. To study this further, we used *RNA velocity* (La Manno *et al*, 2018), a high-dimensional vector that “predicts” the future transcriptional state of individual cells using information of reads mapping to spliced and unspliced transcripts. Applying this to our nascent scRNA-seq data revealed trajectories of proliferating cells towards the senescent state that nicely matched declining *HMGB1* levels (but not the levels of *LMNB1* or *HMGBA1* for example; **Fig. S2E,F**). Hence, the nuclear loss and secretion of HMGB1 is an early and defining event for senescence entry.

### HMGB1 binds active chromatin loci in a cell type-specific manner

Capturing HMGB proteins bound to chromatin has proven challenging, because HMG-box DNA-binding domains are not compatible with standard formaldehyde fixation (Pallier *et al*, 2003; Teves *et al*, 2016). Here, we employed a tailored dual-crosslinking ChIP strategy to efficiently capture HMGB1 bound to its cognate sites genome-wide in both proliferating HUVECs and IMR90 (**Figs 2A** and **S3A**; see **Methods** for details). HMGB1 essentially only binds regions marked by H3K27ac (**Fig. S3B**), the vast majority of which are promoters and gene bodies of active genes (approx. 75% and 65% of 810 and 593 peaks in HUVECs and IMR90, respectively; **Fig. 2B**). Given this, and the fact that HUVECs and IMR90 have very different gene expression programs, their ChIP-seq repertoires only share 40 peaks (**Fig. S3C**). Mapping HMGB1 peaks relative to mononucleosome positioning data from IMR90, we found that although nucleosome-free in proliferating cells, they become occupied upon senescence entry (**Fig. 2C**)

**Figure 2.**
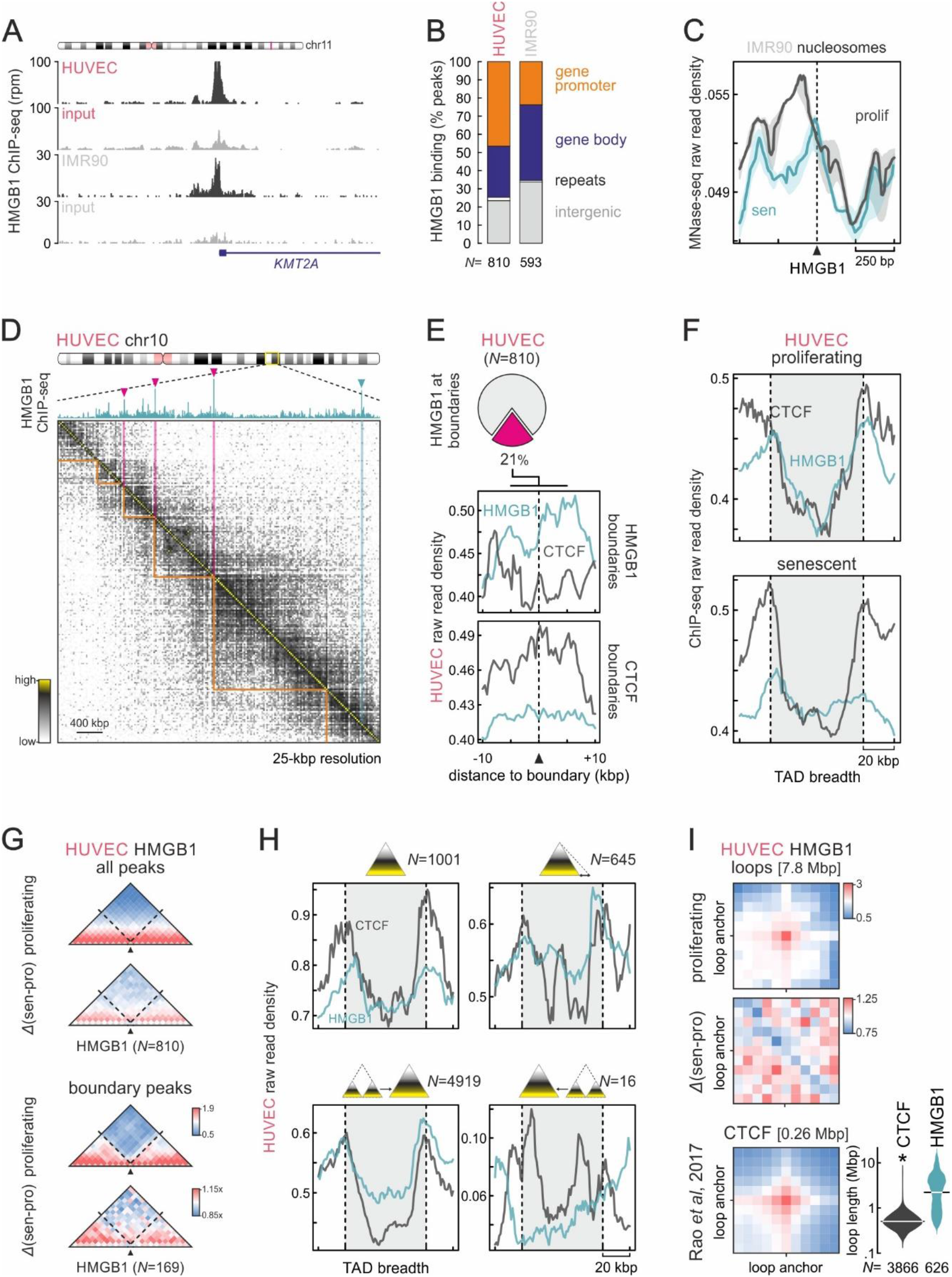
HMGB1 binds active genomic loci and demarcates a subset of TAD boundaries. (A) Genome browser snapshots showing raw HMGB1 ChIP-seq signal at the *KMT2A* promoter in HUVEC and IMR90 (*black);* input tracks (*grey*) provide a negative control. (B) Bar graphs showing the genomic distribution of HMGB1 binding peaks in HUVEC and IMR90. The number of peaks (*N*) analyzed per each cell type is indicated. (C) Line plots showing distribution of mononucleosomal occupancy using MNase-seq data from proliferating (*black*) and senescent IMR90 (*green*) in the 1 kbp around HMGB1 peaks. (D) Exemplary Hi-C heatmap for a subregion of HUVEC chr10 aligned to HMGB1 ChIP-seq; peaks at TAD boundaries (*orange lines*) are indicated (*magenta arrowheads).* (E) Pie chart (*top*) showing the fraction of HUVEC HMGB1 peaks residing at TAD boundaries. Line plots (*below*) showing average HMGB1 (*green*) and CTCF ChIP-seq profiles (*grey*) in the 20 kbp around HMGB1- or CTCF-marked boundaries. (F) Line plots showing average HMGB1 (*green*) and CTCF profiles (*grey*) along TADs ±20 kbp from proliferating (*top*) or senescent HUVECs (*bottom*). (G) Heatmaps showing average Hi-C signal from proliferating or senescent/proliferating HUVEC in the 0.4 Mbp around all HMGB1 peaks (*top*) or those at TAD boundaries (*bottom*). *N* indicates the number of peaks in each subgroup. (H) As in panel F, but for TADs that do not change (*top left*), shift one boundary (*top right*), merge (*bottom left*), or split upon senescence entry (*bottom right). N* indicates the number of TADs in each subgroup. (I) Heatmaps show mean Hi-C signal connecting HMGB1-bound sites from proliferating HUVEC (*top*) and their loss upon senescence entry (*middle);* HUVEC CTCF loops provide a positive control (*bottom*); median CTCF and HMGB1 loop sizes are shown in square brackets and distributions are presented by violin plots. *: significantly different; *P*<0.01; Wilcoxon-Mann-Whitney test.

*De novo* motif discovery in accessible “footprints” of DNase I-hypersensitivity (derived using ENCODE data) within HMGB1 peaks revealed that it binds G/C-rich motifs; however, these do not converge into a single consensus sequence (**Fig. S3D**). We also surveyed these accessible footprints for known transcription factor (TF) motifs to infer co-binding complexes or complexes that might replace HMGB1 on senescent chromatin. We found that HMGB1 binding sites are significantly enriched for E2F-family motifs, which are notably downregulated upon senescence entry. Also, motifs for senescence-activated corepressors (e.g., REST and HEY2) and for the architectural ZBTB7B protein that is important for inflammatory gene activation (Nikopoulou *et al*, 2018) are also enriched therein (**Fig. S3E**). We also noticed that HMGB1 peaks often cluster along chromosomes of proliferating cells (**Fig. S3F**, *top*). This was reminiscent of the clustering of “super-enhancers” (SEs; Hnisz *et al*, 2013), and we applied such an analysis to HMGB1 ChIP-seq peaks. We found that <10% of HMGB1 peaks met the criteria previously set for SE discovery, and these HMGB1 clusters could be linked to 48 genes. However, only few of these were strongly regulated upon senescence entry, with most being upregulated (**Fig. S3F**, *bottom*). This nonetheless agreed with the genome-wide picture where at least twice as many HMGB1-bound genes are up- rather than downregulated in both HUVECs and IMR90. These senescence-induced genes are involved in ECM organization, regulation of apoptosis, as well as in p53 signaling. On the other hand, downregulated genes in both cell types are linked to cell growth and cell cycle regulation (**Fig. S3G**). Together, this data demonstrates how HMGB1 binds specifically to chromatin at loci relevant to the induction of the senescence gene expression program. Strikingly, the majority of HMGB1 target genes respond to its nuclear loss by increasing their transcriptional levels.

### HMGB1 affects a subset of senescence-regulated TAD boundaries

TADs are often considered as the building blocks of chromosomes (Beagan and Phillips-Cremins, 2020), and their boundaries represent genomic sites of strong local insulation for spatial interactions occurring within each TAD. TAD boundaries are often marked by the presence of bound CTCF and/or active gene promoters (Dixon *et al*, 2012; Nora *et al*, 2017). We recently showed that a considerable number of TAD boundaries in proliferating human cells are marked by HMGB2, and that these boundaries are remodeled upon senescence entry (Zirkel *et al*, 2018). We now reasoned that HMGB1 may also function similarly. To test this, we used Hi-C data from proliferating and senescent HUVEC. First, we found that >20% of the 810 HMGB1 peaks reside at TAD boundaries (called at 25-kbp resolution; **Fig. 2D,E**). Remarkably, we identified boundaries marked by HMGB1, but not by CTCF (and the converse; **Fig. 2E**). As would be expected, TADs in senescent chromosomes lose this demarcation (**Fig. 2F**) and HMGB1-marked boundaries exhibit obvious loss of insulation and a reshuffling of spatial interactions across them (**Fig. 2G**). Next, we grouped TADs according to how their boundaries change upon senescence entry. We found that ~1,000 (15%) HUVEC TADs remain invariant in senescence, and their boundaries are more enriched for CTCF than for HMGB1. CTCF and HMGB1 demarcation is comparable in TADs shifting one boundary or collapsing into one larger TAD (~5,500 TADs in total; **Fig. 2H**). Thus, the nuclear loss of HMGB1 in senescence correlates with the remodeling of TADs, but not with one particular trend (as was previously the case for HMGB2; Zirkel *et al*, 2018). These effects also hold true when analyzing HMGB1 and Hi-C data from proliferating and senescent IMR90 (**Fig. S4A-C**).

Finally, we asked whether HMGB1 engages in long-range contact formation. Using HUVEC Hi-C data at its highest (10-kbp) resolution, we found that HMGB1-bound genomic sites give rise to >600 intrachromosomal loops, which collapse upon senescence entry (**Fig. 2I**). These interactions often coincide with the boundaries of TADs or of smaller subTAD domains, but can often extend beyond TAD boundaries to span Mbps (**Figs 2I** and **S4D**). Interestingly, chromatin domains rich in such loops appear depleted of CTCF loops and *vice versa*, and genes at HMGB1 loop anchors often show senescence entry-coupled gene expression changes (**Fig. S4D**). This data suggests that the topological contribution of HMGB1 is important for both genome architecture and gene regulation.

### Spatial TAD co-association reveals functional specialization of chromosome domains

Given that human chromosomes undergo large-scale changes upon replicative senescence entry (Zirkel *et al*, 2018), which are further accentuated in “deep” or oncogene-induced senescence (Criscione *et al*, 2016; Chandra *et al*, 2015), we asked how TADs along each chromosome associate with one another in higher-order “meta-TAD” conformations (Fraser *et al*, 2015). To this end, we employed a bias-free and unsupervised approach inspired from “topologically-intrinsic lexicographic ordering” (TiLO; Johnson, 2014). Here, TADs are treated as nodes in a clustered spatial network tested for robustness via iterative network slicing (**Fig. 3A**; see **Methods** for details). We applied TiLO to TADs derived from proliferating and senescence HUVEC and IMR90 Hi-C data to discover that senescence TAD clusters are larger in size and contain more consecutive TADs on average (**Figs 3B** and **S5A,B**). This is consistent with the general spatial chromatin compaction upon senescence entry (Chandra *et al*, 2015; Criscione *et al*, 2016; Zirkel *et al*, 2018) and the fact that ~75% of HUVEC TADs merge into larger ones (**Fig. 2H**). Closer inspection of clustering results revealed that the statistically significant decrease in standalone (“singular”) TADs (**Fig. 3C**, *arrows)* fed into the increase of TAD clusters containing four or more TADs (**Fig. S5A**). Singular TADs are positioned between multi-TAD clusters (**Figs 3B** and **S5B**). Strikingly, they are enriched for HMGB1-only boundaries compared to the extremities of other TAD clusters (**Figs 3C** and **S5C**). Although both the HMGB1-marked boundaries of singular TADs and those of TAD clusters with >3 TADs showed strong loss of insulation upon senescence entry (**Figs 3D** and **S5D,E**), singular TADs were rich in upregulated genes and more depleted of strongly downregulated ones (**Fig. 3E**). Notably, these two TAD groups harbor most of the genes differentially-regulated upon senescence entry. However, what discriminates them functionally, is the fact that singular TADs uniquely harbour genes associated with the SASP (**Fig. 3F**) and these are induced in an HMGB1-dependent manner upon senescence entry (**Fig. S5F**). Thus, TiLO has the power to identify spatial co-associations of TADs in *cis*, explaining the functional specialization of different genomic domains demarcated by HMGB1.

**Figure 3.**
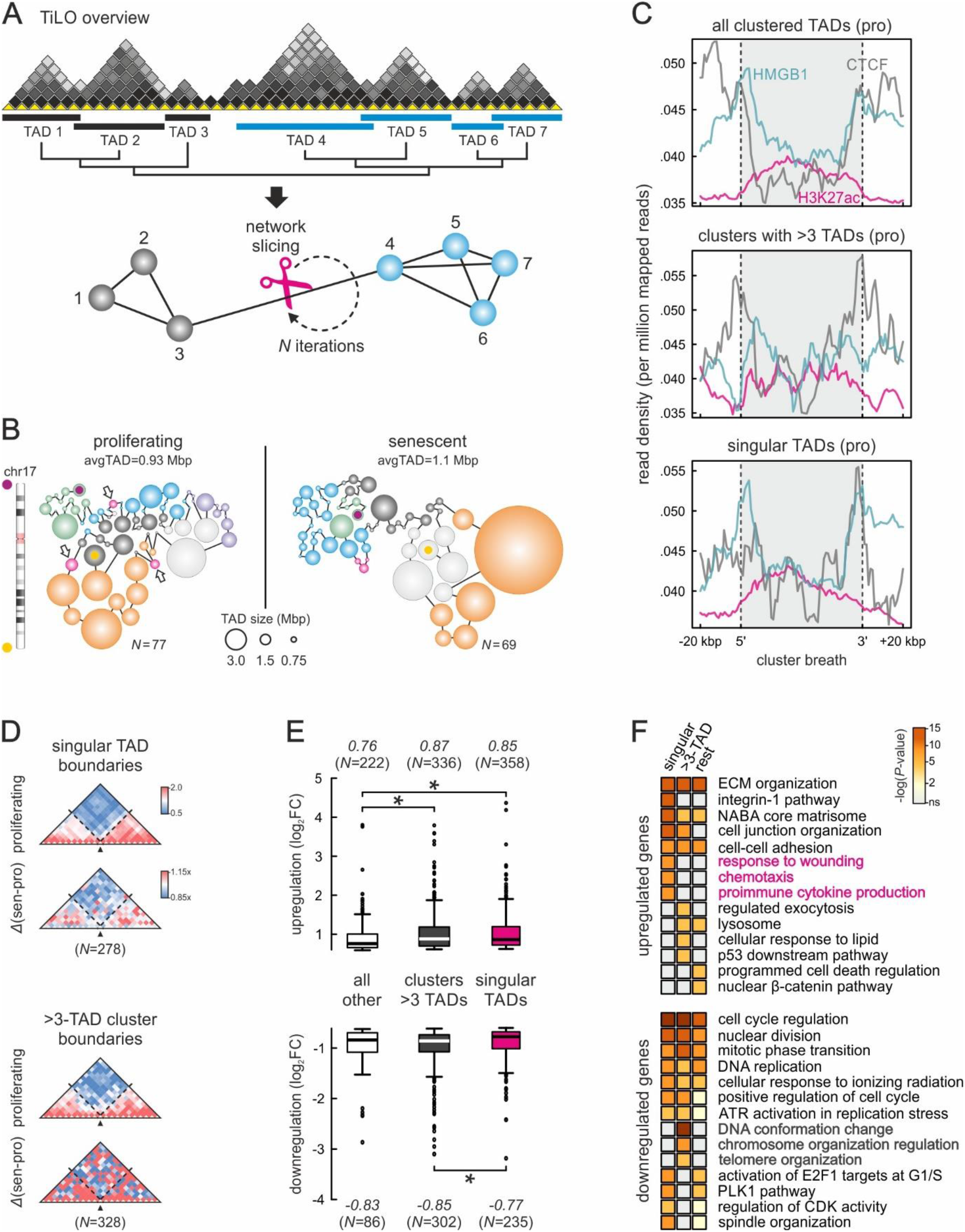
HMGB1 marks individual TADs harboring SASP-related genes. (A) Overview of TiLO. TADs along each chromosome are treated as nodes in an interaction network and inter-TAD Hi-C signal is used to infer network connections. Inferred connections are then sliced and network robustness is assessed iteratively to obtain the final network clustering. (B) Illustration of TAD clusters identified using proliferating (*left*) and senescent HUVEC Hi-C data (*right*) for chr17. Spheres represent TADs; the most 5’/3’ TADs (*purple* and *yellow dots*, respectively) and “singular” TADs (*arrows*) are indicated. (C) Line plots showing average HMGB1 (*green*), CTCF (*grey*), and H3K27ac ChIP-seq profiles (*magenta*) at the extremities of all clustered TADs (±20 kbp; *top*), of clusters of >3 TADs (*middle*) or of “singular” TADs (*bottom*; as those indicated in panel B) from proliferating HUVECs. (D) Heatmaps showing average Hi-C signal from proliferating or senescent/proliferating HUVEC in the 0.4 Mbp around boundaries from singular TADs (*top*) or from clusters of >3 TADs (*bottom*). The number of boundaries in each subgroup (*N*) is indicated. (E) Box plots showing (log_2_) fold-change in mRNA levels of senescence-regulated genes embedded in the three cluster categories from panel C. Median expression fold-change and the numbers of genes in each group are also indicated. *: *P*<0.01; Wilcoxon-Mann-Whitney test. (F) Heatmaps showing GO terms/pathways associated with differentially-expressed genes in the three cluster groups from panel C. Chromatin organization and SASP-related GO terms are highlighted.

### *HMGB1* depletion underlies induction of the senescence transcriptional program

It was previously demonstrated that transduction of WI-38 human fibroblasts with shRNAs against *HMGB1* suffices for senescence induction (Davalos *et al*, 2013). Here, we treated HUVECs with self-delivering siRNAs targeting *HMGB1.* This led to a ~2-fold reduction of HMGB1 protein and RNA levels within 72 h (**Fig. S6A**). It was accompanied by a doubling of β-gal and p21-positive cells in knockdown populations, but by only small changes in nuclear size and BrdU incorporation (**Fig. S6B-E**). To obtain stronger effects, we turned to IMR90 where standard siRNA transfections allowed for a >10-fold decrease in HMGB1 protein and RNA levels, while also regulating expression of known senescence marker genes like *CDKN1B* and *HMGA1* (without affecting *HMGB2* levels; **Fig. 4A**). Analysis of RNA-seq data from siRNA-treated and control IMR90 returned ~900 up- and >950 down-regulated genes upon *HMGB1*-knockdown (**Fig. 5B**). GO term and gene set enrichment analyses showed that the upregulated genes could be linked to the SASP and proinflammatory signalling, while downregulated ones associate with changes in chromatin organization, transcriptional silencing, and the p53 pathway (**Fig. 5C,D**), all hallmarks of senescence entry. Looking for genes that are differentially-regulated upon knockdown but also bound by HMGB1, we identified 44 up- and 56 downregulated genes constituting direct targets. These genes showed enrichment for HMGB1 ChIP-seq signal at their 5’ and 3’ ends, and were linked to NF-κB activation, and to chromatin organization or cell growth signalling, respectively (**Fig. S6F**). Direct comparison of significant gene expression changes upon *HMGB1*-knockdown to those from senescence entry cells revealed poor correlation (**Fig. S6G**, *left*). Genes downregulated in both knockdown and senescent cells were linked to RNA splicing, processing, and cleavage. Interestingly, genes that were downregulated in senescence and upregulated upon *HMGB1*-knockdown were again relevant to RNA metabolism, but also in stress response and apoptosis signalling (**Fig. S6G**, *right).*

**Figure 4.**
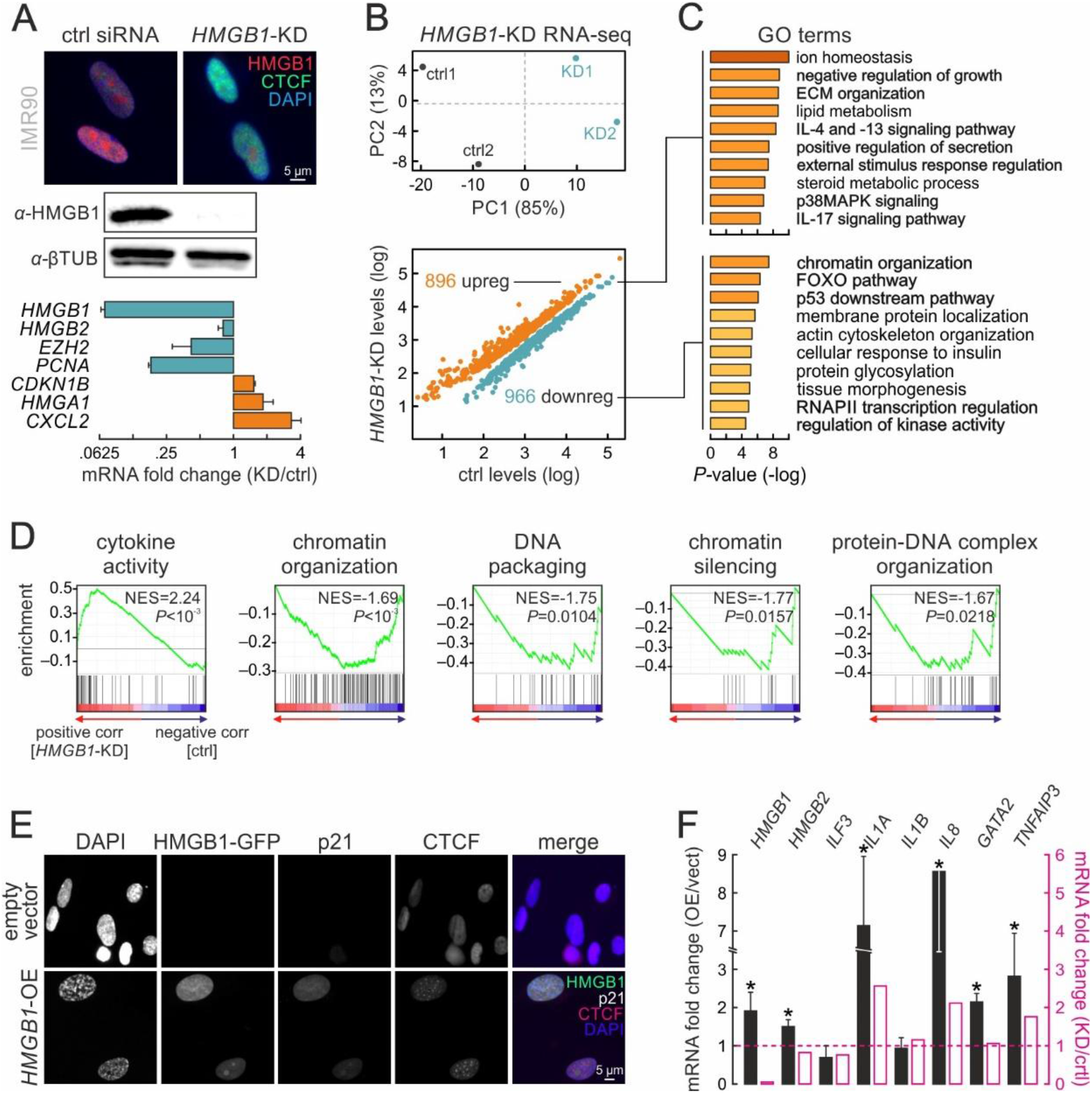
*HMGB1* expression modulation induces senescence-specific gene expression changes. (A) Immunofluorescence (*top*), western blot (*middle*) and RT-qPCR analyses (*bottom*; mean fold-change ±S.D., *N*=3) confirm *HMGB1* knockdown in IMR90. Bar: 5 μm. (B) PCA analysis plot (*top*) of control (*black*) and *HMGB1*-knockdown replicates (*green*). Scatter plot (*bottom*) showing significantly up- (>0.6 log_2_-fold change; *orange*) or downregulated genes (<-0.6 log_2_-fold change; *green*) upon *HMGB1* knockdown. (C) Bar plots showing GO terms associated with the up/downregulated genes from panel B and their enrichment *P*-values (*right).* (D) Gene set enrichment analysis (GSEA) of *HMGB1-KD* data. Normalized enrichment scores (NES) and associated *P*-values for each set are shown. (E) Representative images of IMR90 overexpressing HMGB1-GFP, immunostained for p21 and CTCF, and counterstained with DAPI. IMR90 transfected with empty vectors provide a control. Bar: 5 μm. (F) Bar plots showing RT-qPCR data (mean mRNA fold-change ±S.D., *N*=2) for selected genes in *HMGB1*-overexpressing compared to control IMR90. The mean fold-change for each mRNA from *HMGB1*-knockdown RNA-seq data is also shown for comparison (*magenta bars).* *: *P*<0.01; unpaired two-tailed Student’s t-test.

**Figure 5.**
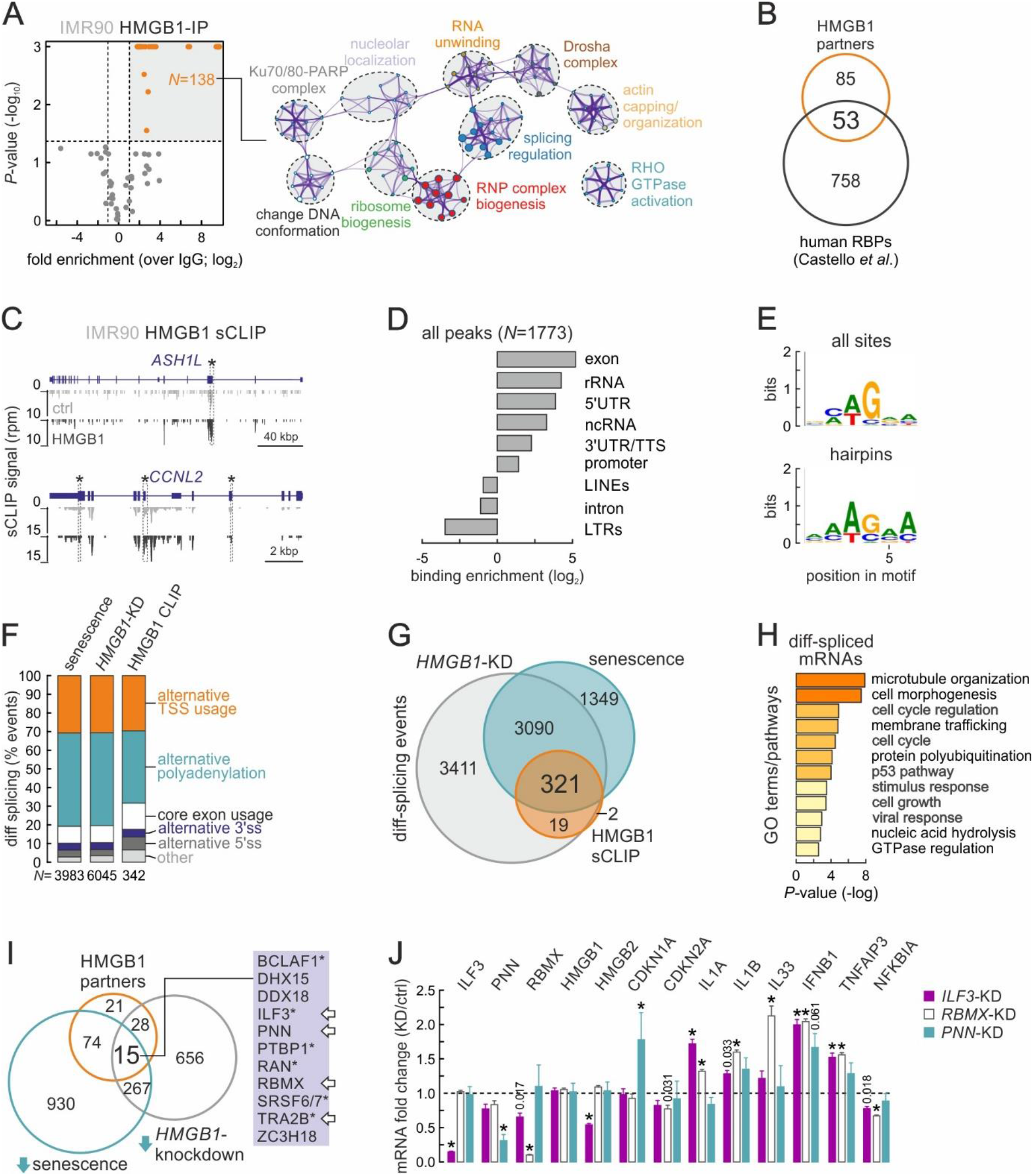
HMGB1 binds specific mRNAs and affects splicing. (A) Volcano plot (*left*) showing mass-spec data for proteins co-immunoprecipitating with HMGB1. Statistically-enriched HMGB1 interactors (*orange dots*) associate with the GO terms/pathways illustrated in the network analysis (*right;* node size reflects the number of proteins it includes; proteins are listed in Table S4). (B) Venn diagram showing 1/3 of HMGB1 interactors classifying as RBPs (according to data by Castello *et al*, 2016). (C) Genome browser views showing HMGB1 sCLIP data (*black*) along the *ASH1L* and *CCNL2* loci; input tracks (*grey*) provide a negative control. *: exemplary significantly-enriched peaks. (D) Bar graphs showing genomic distribution of HMGB1 RNA-bound peaks (log_2_ enrichment). (E) Logos showing consensus motifs deduced via ssHMM analysis from all (*top*) or from hairpin-embedded HMGB1 sCLIP peaks (*bottom).* (F) Bar plots showing relative occurrence of differential-splicing events in IMR90 undergoing senescence (*left*), in HMGB1-knockdown IMR90 (*middle*) or in HMGB1-bound mRNAs (*right).* The number of bound mRNAs (*N*) analyzed is indicated below each bar. (G) Venn diagram showing differential-splicing events shared between conditions from panel F. (H) Heatmaps showing GO terms/pathways associated with differentially-spliced mRNAs shared between the conditions in panel G. (I) Venn diagram (*left*) showing 15 HMGB1 interacting proteins from panel A are also downregulated upon both senescence entry and *HMGB1-knockdown* in IMR90. Of these, 12 are RBPs, 6 have been implicated in senescence (*asterisks*), and 4 are bound by HMGB1 in sCLIP data (*arrows).* (J) Bar graphs showing mean fold-change of selected mRNAs (over ±S.D., *N*=2) from *ILF3-(purple), RBMX-(white*) or *PNN*-knockdown experiments (*green*) in proliferating IMR90. *: significantly different to siRNA controls; *P*<0.01, unpaired two-tailed Student’s t-test.

We complemented our knockdown experiments with *HMGB1* overexpression. We randomly integrated an HMGB1-GFP fusion open reading frame in IMR90 using a doxycycline-inducible piggybac construct (**Fig. 5E**). We selected for transformed cells using antibiotics, but refrained from generating single cell-derived populations in order to gauge heterogeneity due to differences in integration sites. Within <24 h of overexpression induction, nuclear accumulation of HMGB1 in a subset of the population led to a strong increase in p21 signal, to the emergence of characteristic DAPI-dense foci and, in some cases, to the formation of senescence-induced CTCF clusters (Zirkel *et al*, 2018; **Fig. 5E**). All these are hallmarks of replicative senescence entry, and agree with changes in the expression of SASP genes due to *HMGB1* being overexpressed (**Fig. 5F**). Thus, increased HMGB1 titers also suffice for driving entry into senescence, most probably via reinforced and continuous paracrine signalling.

### HMGB1 binds and regulates senescence-related mRNAs independently of its chromatin role

The number of protein-coding loci bound by HMGB1 and regulated upon senescence entry and *HMGB1* knockdown does not explain the full extent of the senescence transcriptional program. To address this disparity, we pursued the idea that HMGB1 also acts as an RNA-binding protein, as was suggested by a recent classification of the human proteome (Castello *et al*, 2016; Trendel *et al*, 2018). This idea was reinforced by our analysis of Ribo- and RNA-seq data (**Figs S1E** and **S6G**), as well as by our cataloguing of the proteins partners of HMGB1 in proliferating IMR90. Mass spectrometry analysis revealed a broad range of RNA-binding proteins and splicing regulators co-immunoprecipitating with HMGB1, in addition to the expected proteins associated with chromatin conformation regulation (**Fig. 5A**). In fact, almost 40% of HMGB1 interactors qualify as RNA-binding proteins (Castello *et al*, 2016; **Fig. 5B**).

To study HMGB1 as a direct RNA binder, we applied sCLIP (Kargapolova *et al*, 2017) to proliferating IMR90 (**Fig. S7A,B**). Analysis of these two well-correlated replicates (**Fig. S7C**) provided a compendium of 1,773 binding peaks on 866 different transcripts, many of which are known to be regulated upon senescence entry (e.g., *ASH1L* and *CCNL2;* **Fig. 5C** and **Table S1**). Reassuringly, HMGB1-bound mRNAs display <25% overlap to HMGB1-bound genes in ChIP-seq. Thus, crosslinking of HMGB1 to RNA is not simply a byproduct of its binding to active loci on chromatin.

On RNA, HMGB1 mostly binds to exons and 5’/3’ UTRs, but also to a substantial number of non-coding RNAs (**Fig. 5D**). HMGB1-bound sites present the same hexameric 5’-NMWGRA-3’ (M=A/C, W=A/T, R=A/G) motif irrespective of the predicted folding of the underlying RNA (**Figs 5E** and **S7D**). Much like what we observed in ChIP-seq, HMGB1 binds ~3-fold more transcripts that are up- rather than downregulated upon senescence (**Fig. S7E**, *left*). Upregulated mRNAs associated with senescence-related GO terms like ECM organization, wound healing and negative regulation of cell proliferation, while downregulated ones mostly with processes like RNA splicing, RNA-/miRNA-mediated gene silencing, or histone remodeling and deacetylation (**Fig. S7E**, *right).* When we crossed sCLIP with RNA-seq data from *HMGB1*-knockdown IMR90, 56 and 97 mRNAs were found to be bound by HMGB1 and up- or downregulated, respectively. Curiously, upregulated transcripts showed a slight bias for HMGB1 binding towards their 5’ ends, while downregulated ones showed evident 3’ end binding (**Fig. S7F**). Consistent with all our previous observations, upregulated mRNAs could be linked to processes like ECM organization and proinflammatory activation, while downregulated mRNAs associated with non-inflammatory signalling and chromatin organization (**Fig. S7G**).

RNA-binding proteins do not only regulate the availability of transcripts, but can also alter their splicing patterns. Thus, we next examined how splicing is altered upon senescence entry by IMR90 using Whippet (Sterne-Weiler *et al*, 2018). We documented ~4,000 significant changes in mRNA splicing, the majority of which concerned the use of alternative transcription start and polyadenylation sites (>80% of cases; **Fig. 5F**). This trend remained essentially invariable when we interrogated the type of splicing changes occurring upon *HMGB1* knockdown or to HMGB1-bound and differentially-spliced mRNAs (**Fig. 5F**). Importantly, the vast majority (>90%) of these splicing events in mRNAs from HMGB1 sCLIP data overlap events seen in both senescent and *HMGB1*-knockdown IMR90 (**Fig. 5G**). Differentially-spliced mRNAs encode factors linked to senescence-regulated processes like cell cycle and cell growth regulation, and the p53 pathway (**Fig. 5H**). Thus, the nuclear loss of HMGB1 directly affects processing of the cell’s transcriptome.

Given this unforeseen role of HMGB1, we revisited its protein interactome (**Fig. 5A**). We found that 15 of the HMGB1 protein partners were downregulated in senescence, but also upon *HMGB1-* knockdown. Of these, 12 qualified as RNA-binding proteins (Castello *et al*, 2016), with BCLAF1 (Shao *et al*, 2017), ILF3 (Wu *et al*, 2015; Tominaga-Yamanaka *et al*, 2012), PTBP1 (Georgilis *et al*, 2018), RAN (Cekan *et al*, 2016; Gu *et al*, 2016; Sobuz *et al*, 2019), SRSF7 (Chen *et al*, 2017), and TRA2B (Chen *et al*, 2018) recently implicated in senescence and SASP regulation (**Fig. 5I**, *starred).* Moreover, the mRNAs of *ILF3, PNN, RBMX* and *TRA2B* are also bound by HMGB1 in IMR90 sCLIP data (**Fig. 5I**, *arrows).* Data in the STRING database (https://string-db.org; Szklarczyk *et al*, 2019) based exclusively on experimental observations link all of them in a tightly woven network relevant to RNA processing (**Fig. S7H**).

To further validate this, we performed co-immunoprecipitation experiments showing that HMGB1 and ILF3 do physically interact (**Fig. S7I**). We further show that although ILF3 is downregulated upon *HMGB1-*knockdown, its proteins levels increase once HMGB1 is overexpressed (**Fig. S7J**). This increase also manifests in a paracrine manner, since cells not carrying the HMGB1-overexpression cassette still show elevated ILF3 levels (**Fig. S7J**, *right).* In addition, the ILF3 increase coincides with NFκB translocation into cell nuclei signifying inflammatory activation of the cells (**Fig. S7K**). In line with what was reported for oncogene-induced senescence (Tominaga-Yamanaka *et al*, 2012), ILF3 binds SASP-relevant mRNAs in proliferating IMR90 (**Fig. S7L**) and its senescence-induced loss will lead to their stabilization for translation. Notably though, in our model, ILF3 also binds HMGB2 transcripts (**Fig. S7L**) and is thus implicated in a feedforward regulatory loop with this factor. We went on to knock *ILF3, RBMX*, and *PNN* down individually in proliferating IMR90. With the exception of p21 induction, *PNN-* knockdown did not affect *HMGB1/2* or SASP-related mRNA levels. In contrast, knocking down *ILF3* or *RBMX* led to the upregulation of interleukins, *IFNB1* or *TNFAIP3* (**Fig. 5J**), all indicative of inflammatory activation. Interestingly, *HMGB1* levels did not change in any of these experimental setups, while *ILF3-* knockdown did suppress *HMGB2.* On this basis, we infer that HMGB1 is an upstream regulator of this whole cascade (given its binding on the mRNAs of all three RBPs; **Fig. 5I**), while ILF3 can specifically modulate HMGB2 (which is not possible via *HMGB1-*knockdown; **Fig. 4A**). Thus, HMGB1 is central to a regulatory circuit comprising RBP cofactors that regulate one another, as well as cell-autonomous and paracrine senescence.

## Discussion

Unlike the well-documented extracellular role of HMGB1 as a proinflammatory stimulus, its positioning along mammalian chromosomes and the gene expression control it exerts are poorly understood. Here, we assign a multifaceted role to HMGB1 — first as an on-chromatin regulator of active gene loci, and then as a *bona fide* RNA-binding protein regulating a distinct subset of mRNAs. Together, we deduce that HMGB1 acts to “buffer” gene expression levels, its loss from senescent cell nuclei mostly triggering upregulation of target loci and mRNAs (**Figs S3G** and **S7E**). Moreover, ~1/5 of HMGB1-bound positions mark TAD boundaries. Many of these TADs harbor SASP-related genes that are induced upon both senescence entry and *HMGB1* knockdown (**Fig. S5**). This suggests that HMGB1 loss also triggers rewiring of chromatin topology linked to gene expression changes. Interestingly, 3D chromatin domains rich in HMGB1 loops are generally depleted of CTCF loops (**Fig. S4D**), and the same applies to a subset of TAD boundaries (**Fig. 2E**). This implies that these spatial conformations might be incompatible and, thus, confer different regulatory modes across chromosomes.

In its novel role as a direct RNA-binding regulator, HMGB1 is part of RNP complexes that affect transcript splicing and processing. In fact, a number of HMGB1 partners are RBPs implicated in the regulation of senescence induction and the SASP (Tominaga-Yamanaka *et al*, 2012; Wu *et al*, 2015; Cekan *et al*, 2016; Gu *et al*, 2016; Shao *et al*, 2017; Chen *et al*, 2017; Chen *et al*, 2018; Georgilis *et al*, 2018; Sobuz *et al*, 2019). In addition, the mRNAs of four of these RBPs (*ILF3, PNN, RBMX* and *TRA2B*) are also direct HMGB1 targets (**Fig. 5I**). This renders HMGB1 a central player also in RNA homeostasis. Once the control is exerts is alleviated due to its depletion from cell nuclei, mRNAs necessary for paracrine senescence become stabilized (Tominaga-Yamanaka *et al*, 2012 and **Fig. S7K**). This constitutes a remarkable example of a regulatory circuit, where programmed deregulation of genes, transcripts and topology in one cellular compartment (the nucleus) is in direct and quantitative crosstalk with signaling deployed in another (in extracellular space via HMGB1 secretion). Our observations come to substantiate previous hypotheses of low nuclear HMGB1 titers being necessary for the full deployment of the SASP (Davalos *et al*, 2013). Thus, the cascade regulating senescence entry has a strong, almost hierarchical, dependency on the nuclear events preceding SASP induction.

Recently, we characterized the function of the sister protein to HMGB1, HMGB2, as regards the entry into replicative senescence (Zirkel *et al*, 2018). The loss of HMGB2 appears to precede that of HMGB1, and drives formation of prominent senescence-induced CTCF clusters (SICCs). This affects the spatial architecture of chromosomes, and concomitantly gene expression. Intriguingly, HMGB2 target loci are also usually upregulated once relieved of HMGB2 binding; however, this is the only similarity between the functions of HMGB1 and HMGB2. The loss of HMGB1 does not trigger SICC formation, the same way that the loss of HMGB2 does not trigger p21 activation or SASP induction. Also, HMGB1 and HMGB2 bind non-overlapping genomic loci and demarcate TADs in distinct modes – HMGB2 marks the extremities of TADs that shift one boundary upon senescence entry, while HMGB1 is mostly found at the boundaries of “singular” TADs that harbour SASP-related genes. Critically, knockdown of *HMGB1* does not reduce *HMGB2* levels in human cells. Conversely, knocking down *HMGB2* does not affect *HMGB1* levels (Zirkel *et al*, 2018), meaning that the pathways these two factors control do not overlap, but are rather deployed in parallel.

Finally, *HMGB1*-knockdown in primary lung fibroblasts leads to gene expression changes that are partially inversed upon senescence entry of the same cells (e.g., the negative regulation of RNAPII transcription is suppressed in the knockdown, but not in senescence; *MYC* activation is upregulated in the knockdown, but suppressed upon senescence entry). This may be interpreted as a coordinated counter-regulation of HMGB1-driven effects on the path to senescence, and can be explained by the fact that the nuclear presence of HMGB1 is linked to favorable proautophagic effects that enhance cell survival and limit programmed cell death (Tang *et al*, 2010). This might also be a simple way to explain the strong overexpression of *HMGB1* in various cancer types (Tang *et al*, 2010; Li *et al*, 2014). This overexpression, although highly deleterious for normal cells, seems to favor increased cell proliferation (Kang *et al*, 2013; Li *et al*, 2014). Thus, the nuclear abundance of HMGB1 (and presumably also of HMGB2) can be seen as a marker for proliferative capacity: senescent cells essentially have no nuclear HMGBs, while continuously dividing cancer cells display levels even higher than those seen in normal tissue. In a next step, deciphering the functional implications behind this “readout” may potentially help us understand how a given cell can escape senescence and acquire a malignant identity.

## Author contributions

SK, AZ, NS, NJ, YK, TG, IP, and AM performed experiments; GL, JK, and AM performed Ribo-seq; MN, NJ, YK, and EGG performed computational analyses; AP and AB analyzed single-cell nascent RNA-seq data; JA and PN provided library preps and high throughput sequencing; AP conceived the study and wrote the manuscript with input from all coauthors.

## Data availability

All sequencing data generated for this study can be found in the NCBI Gene Expression Omnibus (GEO) repository under the accession number GSE98448, with the exception of sCLIP and Ribo-seq data that is available under the accession number GSE146047.

## Acknowledgements

We would like to thank members of all laboratories involved in this study for helpful discussions, and Leo Kurian for critical reading of this manuscript. We thank the CMMC FACS sorting and the CECAD Proteomics facilities for assistance. Work in the lab of AP was supported by CMMC Junior Research Group core funding, by Deutsche Forschungsgemeinschaft grants (PA2456/4-1 and /5-1), by an Else-Kroener-Fresenius-Stiftung “Key project” grant (2015_A125), and by a CMMC grant (C04, with AB). KS was supported by IMPRS-GS funding, and JK by the Irish Research Council (EPSPD/2019/214).

## Methods

### Primary cell culture and senescence markers

HUVECs from single, apparently healthy, donors (passage 2-3; Lonza) were continuously passaged to replicative exhaustion in complete Endopan-2 supplemented with 2% FBS under 5% CO2. Cells were constantly seeded at ~10,000 cells/cm^2^, except for late passages when they were seeded at ~20,000 cells/cm^2^. Single IMR90 isolates (I90-10 and −79, passage 5; Coriell Biorepository) were continuously passaged to replicative exhaustion in MEM (M4655, Sigma-Aldrich) supplemented with non-essential amino acids and 10% FBS under 5% CO2. Senescence-associated β-galactosidase assay (Cell Signaling) was performed according to the manufacturer’s instructions to evaluate the fraction of positively-stained cells at different passages. Cell proliferation was monitored by MTT assays at different passages. In brief, ~5,000 cells are seeded in 96-well format plates in quadruplicates. On the next day, the medium is replaced with 100 ml fresh medium plus 10 ml of a 12 mM MTT stock solution (Invitrogen), and cells are incubated at 37^°^C for 4 h. Subsequently, all but 25 mL of the medium is removed from the wells, and formazan dissolved in 50 mL DMSO, mixed thoroughly and incubated at 37^°^C for 10 min. Samples are then mixed again and absorbance read at 530 nm. Measurements are taken at 24, 48 and 72 h post-seeding, background subtracted, and normalized to the 24 h time point. Finally, nascent DNA synthesis was monitored by EdU incorporation and subsequent labelling with Alexa 488 fluors (Click-iT EdU Imaging Kit; Invitrogen). In brief, cells were incubated in 10 mM EdU for 7 h, fixed using 3.7% PFA/PBS for 15 min at room temperature, permeabilized, and labelled as per manufacturer’s instructions, before imaging on a widefield Leica microscope.

### Immunofluorescence and image analysis

Proliferating and senescent cells were grown on coverslips from the stage indicated and were fixed in 4% PFA/PBS for 15 min at room temperature. After washing once in PBS, cells were permeabilized in 0.5% Triton-X/PBS for 5 min at room temperature. Blocking with 1% BSA/PBS for 1h was followed by incubation with the following primary antibodies for 1-2 h at the indicated dilution: mouse monoclonal anti-HMGB1 (1:1000; Abcam ab190377-1F3); rabbit polyclonal anti-HMGB2 (1:1000; Abcam ab67282); mouse monoclonal anti-HMGB1/2 (1:1000; Sigma-Aldrich 12248-3D2); rabbit polyclonal anti-CTCF (1:500; Active motif 61311); rabbit polyclonal anti-H3K27me3 (1:1000; Diagenode C15410069); mouse monoclonal anti-p21 (1:500; Abcam ab184640-GT1032); rabbit polyclonal anti-lamin B1 (1:2000; Abcam ab16048); mouse monoclonal anti-β-tubulin (1:1000; Sigma-Aldrich T0198-D66). Following immunodetection, cells were washed twice with PBS for 5 min before incubating with secondary antibodies for 1 h at room temperature. Nuclei were stained with DAPI (Sigma-Aldrich) for 5 min, washed, and coverslips mounted onto slides in Prolong Gold Antifade (Invitrogen). Note that for gSTED microscopy only, the 2C Pack STED 775 secondary antibodies (1:2000; Abberior 2-0032-052-6) were used. For image acquisition, a widefield Leica DMI 6000B with an HCX PL APO 63x/1.40 (Oil) objective was used; confocal and super-resolution images were acquired on a Leica TCS SP8 gSTED microscope with a 100x/1.40 (Oil) STED Orange objective. For immunofluorescence image analysis, the NuclearParticleDetector2D of the MiToBo plugin (ver. 1.4.3; available at http://mitobo.informatik.uni-halle.de/index.php/Main_Page) was used. Measurements of nuclear immunofluorescence signal were automatically generated using a mask drawn on DAPI staining to define nuclear bounds. Background subtractions were then implemented to precisely determine the mean intensity per area of each immunodetected protein. Deconvolution of super-resolution images was performed using the default settings of the Huygens software (Scientific Volume Imaging).

### Whole-cell protein extraction, western blotting, and mass spectrometry

For assessing protein abundance at different passages, ~4 x 10^6^ cells were gently scraped off 15-cm dishes, and pelleted for 5 min at 600 x g. The supernatant was discarded, and the pellet resuspended in 100 mL of ice-cold RIPA lysis buffer (20 mM Tris-HCl pH 7.5, 150 mM NaCl, 1 mM EDTA pH 8.0, 1 mM EGTA pH 8.0, 1% NP-40, 1% sodium deoxycholate) containing 1x protease inhibitor cocktail (Roche), incubated for 20 min on ice, and centrifuged for 15 min at >20,000 x g to pellet cell debris and collect the supernatant. The concentration of the nuclear extracts was determined using the Pierce BCA Protein Assay Kit (Thermo Fisher Scientific), before extracts were aliquoted and stored at −70^°^C to be used for western blotting. Resolved proteins were detected using the antisera mentioned above, plus a mouse monoclonal anti-H3K9me3 (1:200; Active motif 39286). For whole-cell proteomics, extracts in RIPA buffer were analyzed by the CECAD proteomics core facility in biological triplicates.

### Chromatin immunoprecipitation (ChIP) sequencing and analysis

For each batch of ChIP experiments ~25 million proliferating cells, cultured to > 80% confluence in 15cm dishes, were crosslinked in 1.5 mM EGS/PBS (ethylene-glycol-bis-succinimidyl-succinate; Thermo) for 20 min at room temperature, followed by fixation for 40 min at 4^°^C in 1%PFA. From this point onward, cells were processed via the ChIP-IT High Sensitivity kit (Active motif) as per manufacturer’s instructions. In brief, chromatin was sheared to 200-500 bp fragments on a Bioruptor Plus (Diagenode; 2x 9 cycles of 30 sec *on* and 30 sec *off* at the highest power setting), and immunoprecipitation was carried out by adding 4 mg of a monoclonal HMGB1 antiserum (Developmental Studies Hybridoma Bank; PCRP-HMGB1-4F10) to ~30 mg of chromatin and rotating overnight at 4^°^C in the presence of protease inhibitors. Following addition of protein A/G agarose beads and washing, DNA was purified using the ChIP DNA Clean & Concentrator kit (Zymo Research) and used in next-generation sequencing on a HiSeq4000 platform (Illumina) to obtain at least 25 million reads were obtained of both sample and its respective “input”. Raw reads (typically 100 bp-long) were mapped to the reference human genome (hg19) using BWA (Li and Durbin, 2010), and the resulting .BAM files were processed using Picard tools (http://broadinstitute.github.io/picard/) before MACS2 software (Zhang *et al*, 2008) was used to identify signal enrichment over input. Thresholded HMGB1 ChIP-seq peaks per each cell type were annotated using Chipseeker (Yu *et al*, 2015) and are listed in **Table S2**; .BAM files were used in *ngs.plot* (Shen *et al*, 2014) for plotting signal coverage over particular genomic positions for different conditions/cell types. Finally, transcription factor recognition motif enrichments within DHS footprints under HMGB1 ChIP-seq peaks were calculated using the Regulatory Genomics Toolbox (Gusmao *et al*, 2014). Note that all other ChIP-seq datasets used here come from previous work (Zirkel *et al*, 2018).

### Whole-genome chromosome conformation capture (Hi-C) and TiLO analysis

Hi-C data from proliferating and senescent HUVEC were generated previously (Zirkel *et al*, 2018), and the HiTC Bioconductor package was used to annotate, correct data for biases in genomic features (Servant *et al*, 2012), and visualize 2D heatmaps with a maximum resolution of 20-kbp at which TADs were also called via TADtool (Kruse *et al*, 2016). For plotting insulation and “loop-o-gram” heatmaps, normalized interactions values in the twenty 20-kbp bins around each HMGB1 peak were added up, normalized to the median value in each matrix and plotted provided the local maxima are higher than the third quantile of Hi-C data in the matrix. All R scripts were described previously (Zirkel *et al*, 2018). HMGB1-anchored loops are listed in **Table S2**.

For Topologically-intrinsic Lexicographic Ordering (TiLO), we directly applied an algorithm from mathematical knot theory that makes zero assumptions about the structure, shape or number of clusters in the data (Johnson, 2014). In brief, topologically-intrinsic ordering was used to permutate the linear order of TADs (as a starting organization level in the Hi-C matrices) until a certain “robustly irreducible” topological condition is satisfied. Then, the “pinch ratio” algorithm is used (Heisterkamp and Johnson, 2013) and applied to heuristically slice the network at connections between TADs where local interaction minima are, while also considering noise in the matrices. Finally, this analysis returns a list of TADs grouped into multiple clusters in *cis*, also via its built-in measure for network robustness defining the end-point.

### Single-cell nascent RNA sequencing and analysis

Proliferating (p. 4) and senescent HUVEC (p. 16) were washed once in an isotonic near-physiological buffer (PB) that maintains the cells’ transcriptional activity and subjected immediately to the first steps of the “factory RNA-seq” protocol (Melnik *et al*, 2016). In more detail, cell nuclei are gently isolated using PB+0.4% NP-40, DNase I-treated at 33^°^C for 25 min to detach most chromatin, pelleted and washed once in ice-cold PB, before polyadenylation of nascent RNA as described (Kargapolova *et al*, 2017). Next, ~2,500 cells from each state were loaded onto the Chromium 10X Genomics platform for encapsulation in oil droplets and generation of barcoded cDNA libraries from individual nuclei as per manufacturer’s instructions. Despite the documented 0.8% chance of capturing a cell duplet on this platform, HUVEC nuclei are particularly prone to aggregation. As a result, 494 proliferating and 129 senescent single nuclei were efficiently captured and processed. Following sequencing on a HiSeq4000 platform (Illumina), and mapping to the reference genome (hg38) using STAR (Dobin *et al*, 2013) and filtering via UMI-tools (Smith *et al*, 2017), ~45,000 and ~60,000 reads were generated per each proliferating or senescent cell, respectively. Poor quality cells were excluded (i.e. cells with <300 or >5,000 expressed genes), as were genes expressed <10 cells. This returned 1,650 robustly captured transcripts per cell on average, with >55% of reads mapping to introns or exon-intron junctions, 575 genes being expressed in at least 25% of all cells, and with the 50 most-expressed genes taking over ~22% of all sequencing reads. Mapped and filtered data were then processed and visualized using a compilation of ZINB-WaVE (ver. 1.3.4; Risso *et al*, 2018) and Seurat (ver. 2.3.4; Butler *et al*, 2018) for clustering and t-SNE visualization. ZINB-WaVE was used to create a low-dimensional representation of the gene expression matrix, and factors inferred in the ZINB-WaVE model were added as one of the low-dimensional data representations in the Seurat object in order to identify cell subpopulations via a shared nearest neighbour (SNN) modularity optimization-based clustering algorithm as applied in the *FindClusters* function of Seurat. Visualization was performed using the t-SNE map by applying the *Rtsne* function on the ZINB-WaVE output. This map was integrated into the Seurat object and used to plot gene expression. For differential gene expression analysis between the clusters, we applied a combination of ZINB-WaVE and DESeq2 (Love *et al*, 2014), where the posterior probabilities that counts were generated from the negative binomial count component of the ZINB-WaVE model were used as observation-level weights in the estimation of regression parameters in DESeq2 (Van den Berge *et al*, 2018). Differentially-expressed genes identified via this method were filtered using a threshold of log_2_FC>±1 plus *P*-value<0.05 and are listed in **Table S3**.

### HMGB1 sCLIP and analysis

sCLIP was performed on ~25 million UV-crosslinked nuclei from proliferating IMR90 as previously described (Kargapolova *et al*, 2017) using the same the monoclonal HMGB1 antiserum (DSHB; PCRP-HMGB1-4F10) as for ChIP. Following sequencing of strand-specific libraries on a HiSeq4000 platform (Illumina), raw reads were mapped to the human reference genome (hg19). Consistent peaks were identified by overlapping intervals of peaks with a *P*-value <0.05 from 2 biological replicates to obtain 1773 peaks. This peak annotation was used to count reads uniquely aligned to each peak region using HTSeq, HMGB1-bound transcript coordinates were retrieved via Ensembl (GRCh37) and annotated using HOMER (http://homer.ucsd.edu), and Gene Ontology analysis was performed using Metascape (www.metascape.org). Finally, the final merged peak list was use for *de novo* motif analysis using ssHMM (Heller *et al*, 2017) and significantly enriched motifs were next compared to existing RBP motifs to predict proteins potentially recognizing similar sequences using Tomtom (http://meme-suite.org/tools/tomtom). All HMGB1-bound mRNAs are listed in **Table S1**.

### siRNA-mediated *HMGB1* knockdown

HUVECs were seeded at ~20,000 cells/cm^2^ the day before transfection. Self-delivering Accell-siRNA pools (Dharmacon) targeting *HMGB1*, plus a non-targeting control (NTC; fluorescently-tagged to allow transfection efficiency to be monitored), were added to the cells at a final concentration of 1 mM. Knockdown efficiency was assessed 72 h after transfection using RT-qPCR and immunofluorescence. For IMR90 cells, transfections using standard siRNAs and RNAiMAX (Invitrogen) were carried out as previously described (Zirkel *et al*, 2018).

### Total RNA isolation, sequencing, and analysis

Control and *HMGB1*-knockdown were harvested in Trizol LS (Life Technologies) and total RNA was isolated and DNase I-treated using the DirectZol RNA miniprep kit (Zymo Research). Following selection on poly(dT) beads, barcoded cDNA libraries were generated using the TruSeq RNA library kit (Illumina) and were paired-end sequenced to at least 50 million read pairs on a HiSeq4000 platform (Illumina). Raw reads were mapped to the human reference genome (hg19) using default settings of the STAR aligner (Dobin *et al*, 2013), followed by quantification of unique counts using *featureCounts* (Liao *et al*, 2014). Counts were further normalized via the RUVs function of RUVseq (Risso *et al*, 2014) to estimate factors of unwanted variation using those genes in the replicates for which the covariates of interest remain constant and correct for unwanted variation, before differential gene expression was estimated using DESeq2 (Love *et al*, 2014). Genes with an FDR <0.01 and an absolute (log_2_) fold-change of >0.6 were deemed as differentially-expressed and listed in **Table S3**. For splicing analysis, a reference index on the basis of hg19 annotation was first constructed, combined with all splice sites contained in the mapped RNA-seq reads. Raw reads were then aligned using Whippet (Sterne-Weiler *et al*, 2018) to the constructed index in order to quantify and annotate alternative splicing events. Subsequent plots were plotted using BoxPlotR (http://shiny.chemgrid.org/boxplotr/) and GO term enrichment bar plots using Metascape (http://metascape.org/gp/index.html; Zhou *et al*, 2019).

### Ribo-seq and analysis

High throughput ribosome profiling (Ribo-seq) on proliferating and senescent IMR90 was performed in collaboration with Ribomaps Ltd (https://ribomaps.com) according to an established protocol (Ivanov *et al*, 2018). Three independent replicas of proliferating or senescent IMR90 were grown, harvested in ice-cold polysome isolation buffer supplemented with cyclohexamide, and shipped to Ribomaps for further processing and library preparation. Approx. 15% of each lysate was kept for isolation of RNA and used for RNA-seq of poly(A)-enriched fractions on a HiSeq2500 platform (Illumina). Following sequencing of both Ribo- and mRNA-seq libraries, the per base sequencing quality of each replicate passed the quality threshold, raw read counts were assigned to each protein-coding open reading frame (CDS) for Ribo-seq and to each transcript for mRNA-seq, and replicate correlations were tested.

Read length distribution for Ribo-seq datasets fell within the expected range of 25-35 nt, with a peak between 28-32 nt showing strong periodic signals and an enrichment in annotated CDSs. For mRNA-seq, read lengths ranged between 47-51 nt and distributed uniformly across transcripts. For differential gene expression analysis, Anota2seq (Oertlin *et al*, 2019) was used. Changes in Ribo-seq data represent changes in the ribosome occupancy of the annotated protein coding open reading frame (CDS) and, thus, only ribosome-protected fragments that map to the CDS were used in the analysis. VST normalized counts outputted using DESeq2 (Love *et al*, 2014) and inputted into anota2seq were used for all subsequent downstream analysis. Differences on genes that pass a default false discovery rate (FDR) threshold of 15% were considered regulated. Such significant differences are then categorized into one of the following three modes: (i) translational: changes in Ribo-seq that are not explained by changes in RNA-seq and imply changes at the protein level are due to changes at the translational level; (ii) mRNA abundance: matching changes in RNA-Seq and Ribo-Seq that infer changes at the protein level are predominantly induced by changes at the transcriptional level; (iii) buffering: changes in RNA-seq that are not explained by changes in Ribo-seq and suggest maintenance of constant protein levels induced by changes at the transcriptional level or *vice versa.* Detailed results per gene locus and condition are listed in **Table S5**.

### Statistical tests

*P*-values associated with Student’s t-tests, Fischer’s exact tests or Chi-square Goodness-of-Fit tests were calculated via GraphPad (https://graphpad.com/), and those associated with the Wilcoxon-Mann-Whitney test via U test (https://www.socscistatistics.com/tests/mannwhitney/Default2.aspx). Unless otherwise stated, *P-*values <0.01 were deemed as statistically significant.

**Supplemental Material** contains Figures S1-S7 and Tables S1-S5.

