## Supplemental Figures S1-S7 and Tables S1-S5 for "HMGB1 coordinates SASP-related chromatin folding and RNA homeostasis on the path to senescence"

This **Supplemental Material** contains **Figures S1-S7** and **Tables S1-S5**.

### Supplemental Figures

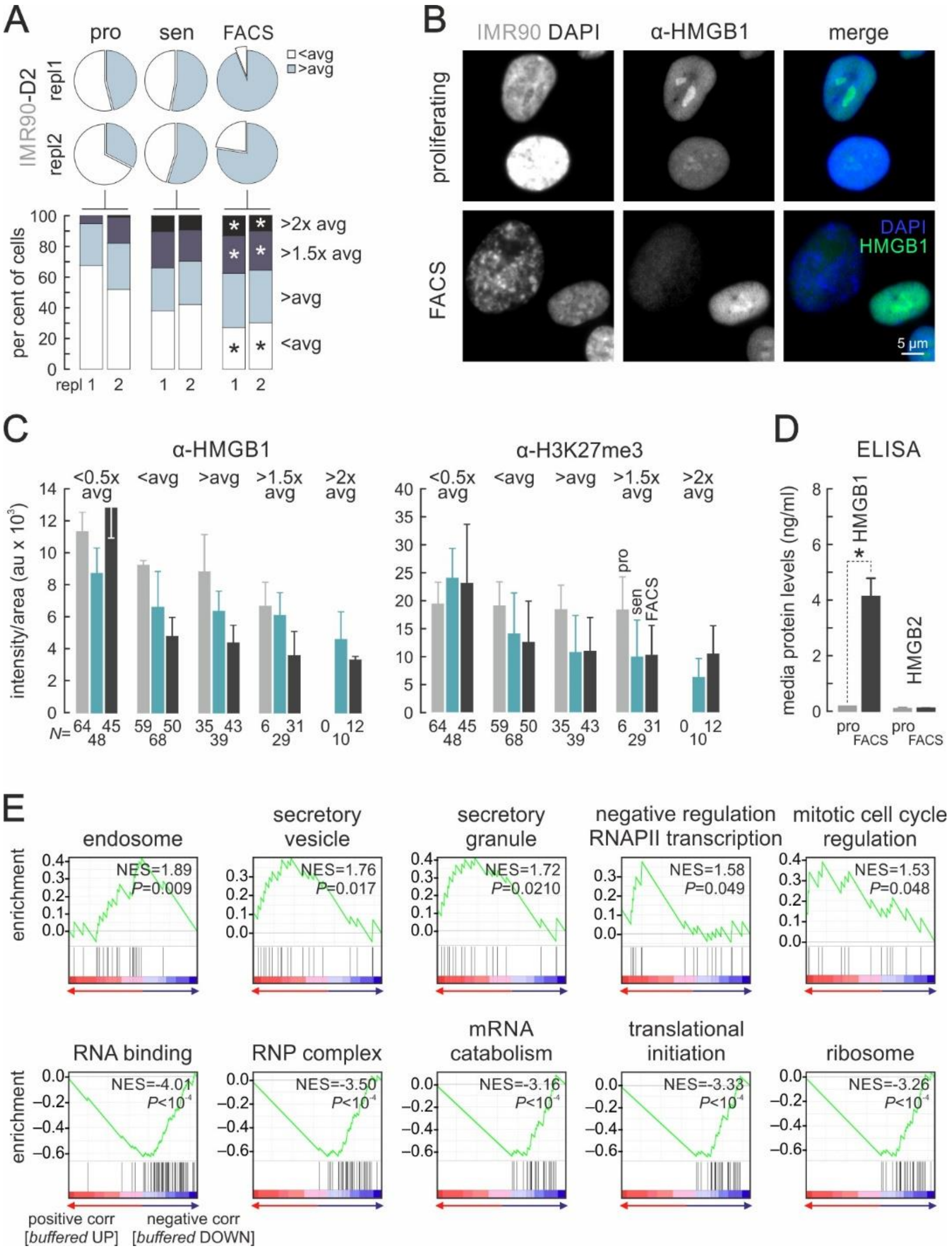


**Figure S1. HMGB1 depletion occurs in enlarged nuclei and is marked by gene expression changes.**

(A) Pie charts (*top*) showing senescent IMR90 populations containing larger-than-average cell nuclei, which can be enriched via FACS. Bar graphs (*bottom*) show stratification of cells according to increasing nuclear size (compared to the population average). *: significantly different to proliferating cells, *P*<0.05; Fisher’s exact test.

(B) Representative images of proliferating (*top row*) and FACS-sorted senescent IMR90 (*bottom row*) immunostained for HMGB1 and counterstained by DAPI. Bar: 5 µm.

(C) Bar graphs showing declining HMGB1 (*left*) and H3K27me3 levels (mean ± S.D.; *right*) in proliferating (*grey*), senescent (*green*) and FACS-sorted IMR90 (*black*) stratified according to nuclear size from images like those in panel B. The number of cells analyzed in each subgroup (*N*) is indicated.

(D) Bar graphs showing HMGB1/B2 levels (mean ± S.D.) detected in the growth media of proliferating (*grey*) or FACS-sorted IMR90 (*black*). *: $P<0.01$; unpaired two-tailed Student's t-test ($N=2$).

(E) Gene set enrichment analysis (GSEA) of genes whose expression is "buffered" in Ribo-seq data. Normalized enrichment scores (NES) and associated *P*-values for each set are shown.

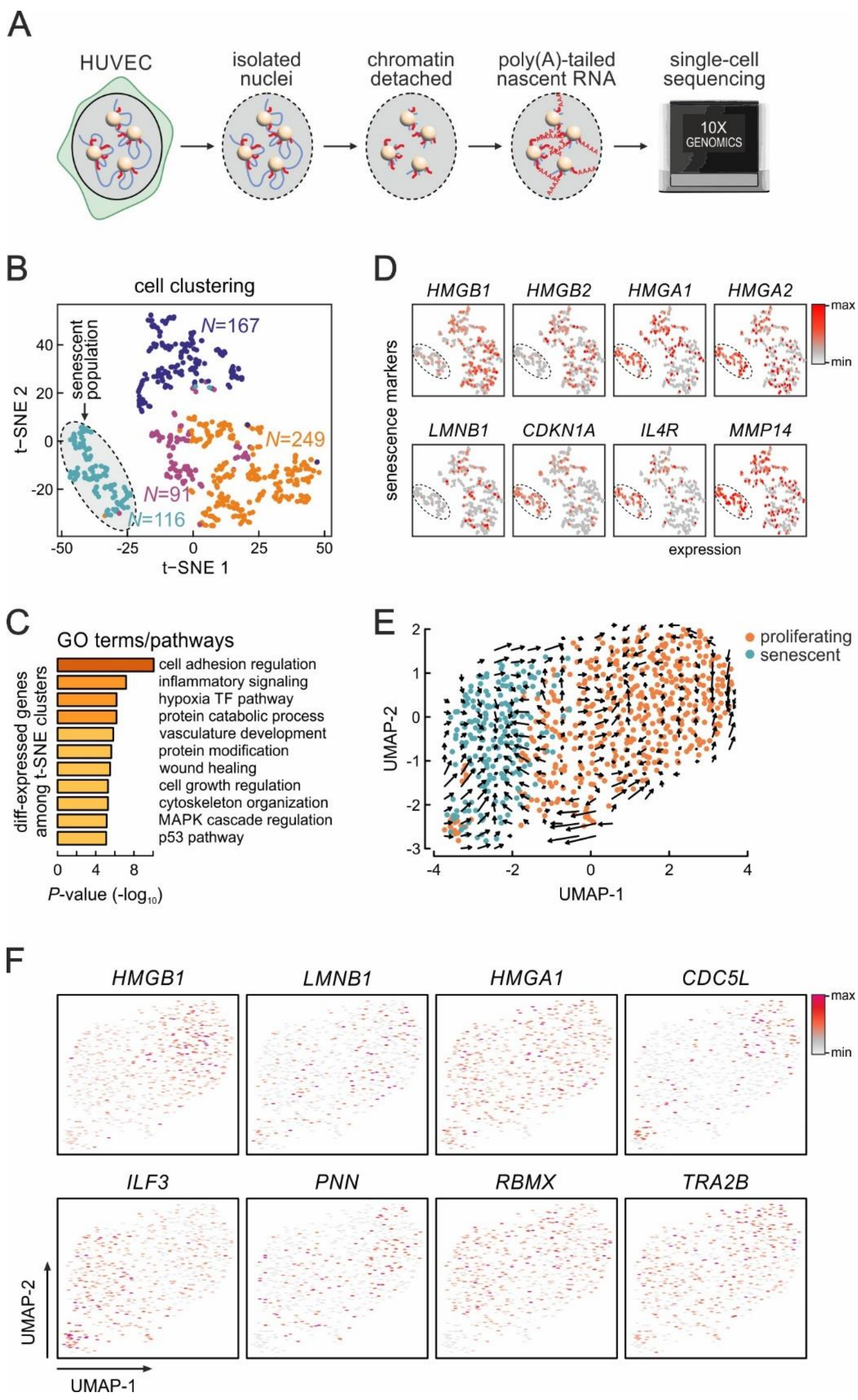


### Figure S2. Single-cell sequencing of nascent RNA.

(A) Strategy for nascent scRNA-seq involving isolation of intact HUVEC nuclei, detachment of non-transcribed chromatin (*blue*) via DNase I digestion, and *in situ* polyadenylation of nascent RNA (*red*) at transcription foci (*spheres*), before standard processing on a 10X Genomics platform.

(B) t-SNE clustering of data from 623 nuclei separates three proliferating cell groups (*blue*, *red*, *purple*, and *green*) from the senescent ones (*grey oval*); *N* indicates the number of cells per cluster.

(C) Bar graphs showing (-log) enrichment *P*-values for GO terms associated with differentially-expressed genes amongst the clusters from panel B.

(D) Heatmaps showing normalized expression levels of selected senescence marker genes in individual cells clustered as in panel B.

(E) UMAP reduction plot showING results from *Velocity* analysis. Arrows indicate trajectories of proliferating HUVECs (*orange*) towards senescent ones (*green*) based on unspliced/spliced RNA ratios in individual cells.

(F) Heatmaps showing normalized expression levels of selected genes in individual cells clustered as in panel E.

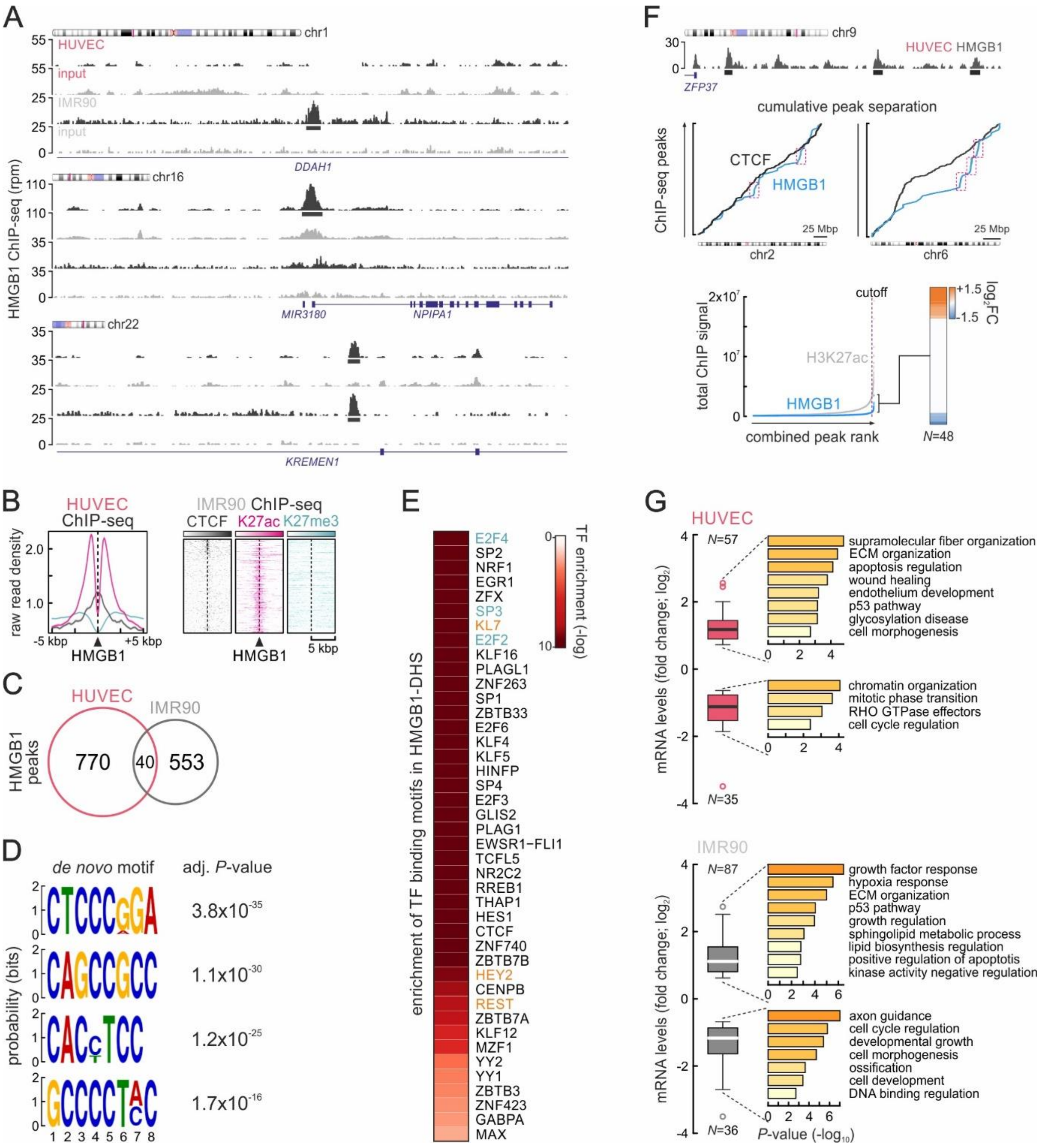


**Figure S3. HMGB1 chromatin-binding features in proliferating cells.**

(A) Genome browser views showing raw HMGB1 ChIP-seq signal at loci in chr1, 16, and 22 from both HUVEC and IMR90 (*black*); input tracks (*grey*) provide a control.

(B) Line plots and heat maps showing distribution of H3K27ac (*magenta*), H3K27me3 (*light blue*), and CTCF ChIP-seq signal (*grey*) in the 20 kbp around HMGB1 peaks from HUVECs and IMR90, respectively.

(C) Venn diagrams showing minimal overlap between HMGB1 binding peaks from HUVEC and IMR90. Overlap is not more than what is expected by chance; $P>0.1$, Chi-square Goodness-of-Fit test.

(D) Logos and associated adjusted *P*-values for the most enriched *de novo* motifs discovered in DNase I-accessible footprints under HMGB1 peaks.

(E) Heatmap showing most enriched TF binding motifs in the footprints from panel C; up- (>0.6 $\log_2$-fold change; *orange*) or downregulated TFs (<-0.6 $\log_2$-fold change; *green*) are indicated.

(F) Genome browser views (*top*) showing clustered HMGB1 ChIP-seq peaks (*black boxes*) near the ZFP37 gene promoter on HUVEC chr9. Plots (*middle*) showing the cumulative distribution of HMGB1 ChIP-seq peaks along HUVEC chr2, 6 and 9 (*green*); CTCF peak distributions (*black*) provide a control. Plot (*bottom left*) ranking "stitched" H3K27ac-marked enhancers (*grey*) and HMGB1-bound sites; those over the cutoff (*magenta*) qualify as "superenhancers". mRNA fold-changes ($\log_2$) of the 48 genes linked to HMGB1 "superenhancers" are plotted as a heatmap (*bottom right*).

(G) Box plots showing mRNA fold-changes ($\log_2$) of genes differentially-expressed upon senescence and bound by HMGB1 in HUVECs (*top*) and IMR90 (*bottom*). GO terms associated with each subgroup and their enrichment *P*-values are shown. The number of genes in each group (*N*) is indicated.

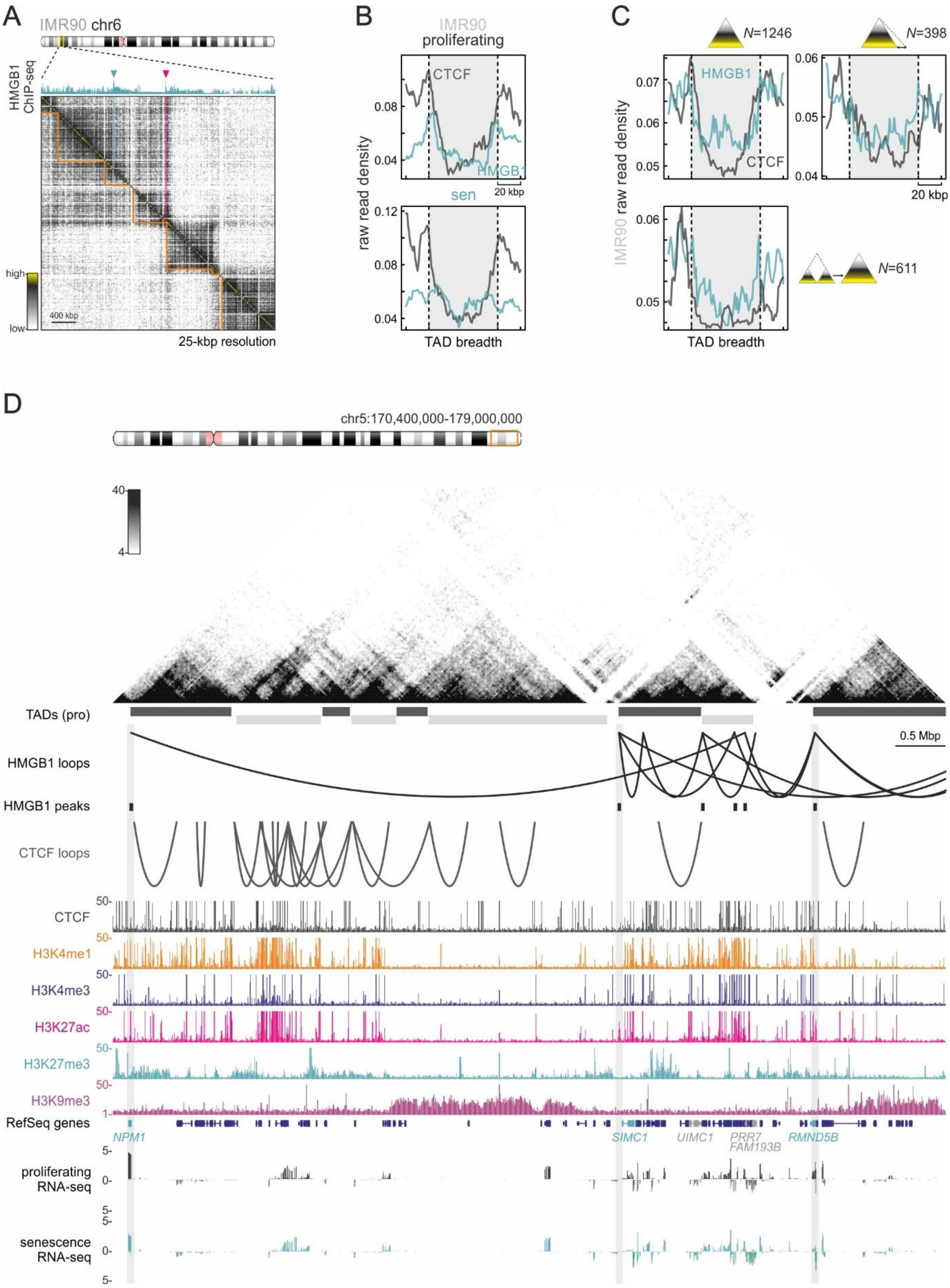


**Figure S4. HMGB1 demarcates TAD boundaries in proliferating IMR90 and HUVECs.**

(A) Exemplary Hi-C heatmap for a subregion in IMR90 chr6 aligned to HMGB1 ChIP-seq; peaks at TAD boundaries (*orange lines*) are indicated (*magenta arrowheads*).

(B) Line plots showing average HMGB1 (*green*) and CTCF ChIP-seq profiles (*grey*) along TADs ±20 kbp from proliferating (*top*) or senescent IMR90 (*bottom*).
(C) As in panel F, but for TADs that do not change (*top left*), shift one boundary (*top right*) or merge upon senescence entry (*bottom left*). *N* indicates the number of TADs in each subgroup.
(D) Exemplary Hi-C data (25-kbp resolution) in an 8.6-Mbp region of HUVEC chr5 aligned to positions of TADs, HMGB1 and CTCF loops, as well as to ENCODE ChIP-seq and own RNA-seq data. The positions of HMGB1 peaks at downregulated promoters (*green gene names*) are highlighted (*grey*).

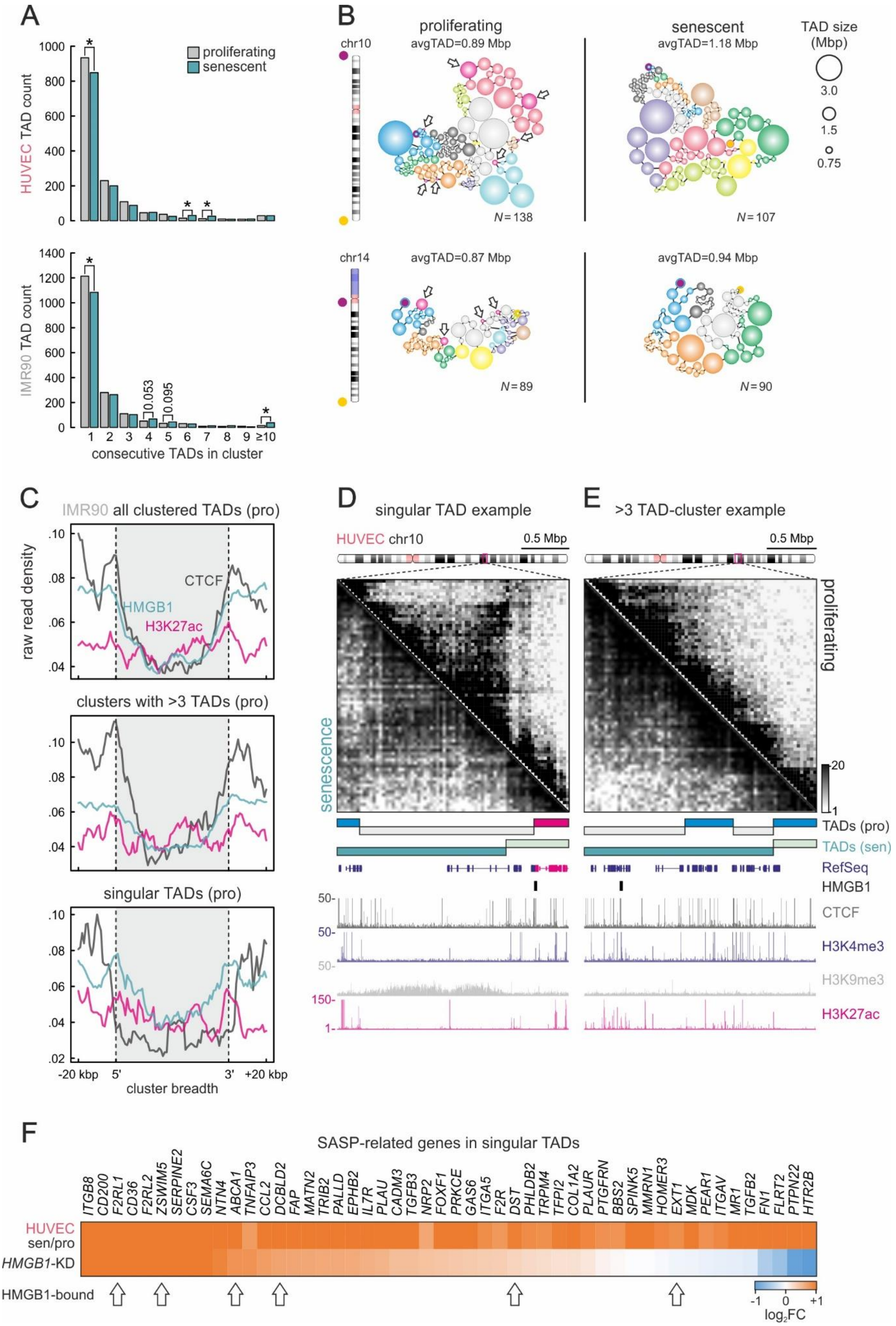
A
HUVEC TAD count
proliferating
senescent
IMR90 TAD count
0.053
0.095
consecutive TADs in cluster
B
chr10
proliferating
avgTAD=0.89 Mbp
N=138
senescent
avgTAD=1.18 Mbp
N=107
TAD size (Mbp)
3.0
1.5
0.75
chr14
avgTAD=0.87 Mbp
N=89
avgTAD=0.94 Mbp
N=90
C
IMR90 all clustered TADs (pro)
raw read density
CTCF
HMGB1
H3K27ac
clusters with >3 TADs (pro)
singular TADs (pro)
-20 kbp
5'
3'
+20 kbp
cluster breadth
D
singular TAD example
HUVEC chr10
0.5 Mbp
senescence
E
>3 TAD-cluster example
0.5 Mbp
proliferating
TADs (pro)
TADs (sen)
RefSeq
HMGB1
CTCF
H3K4me3
H3K9me3
H3K27ac
F
SASP-related genes in singular TADs
ITGB8 CD200 F2RL1 CD36 F2RL2 ZSWIM5 SERPINE2 CSF3 SEMA6C NTN4 ABCA1 TNFAIP3 CCL2 DCBLD2 FAP MATN2 TRIB2 PALLD EPHB2 IL7R PLAU CADM3 TGFB3 NRP2 FOXF1 PRKCE GAS6 ITGA5 F2R DST PHLDB2 TRPM4 TFPI2 COL1A2 PLAUR PTGFRN BBS2 SPINK5 MMRN1 HOMER3 EXT1 MDK PEAR1 ITGAV MR1 TGFB2 FN1 FLRT2 PTPN22 HTR2B
HUVEC sen/pro
HMGB1-KD
HMGB1-bound
log2FC

Figure S5. HMGB1 demarcates specific TAD subsets in primary human cells.

(A) Bar plots showing the number of TADs contained in progressively larger clusters derived using TiLO on proliferating (*grey*) and senescent Hi-C data (*green*) from HUVEC (*top*) and IMR90 (*bottom*). *: $P$<0.01; Fisher's exact test.

(B) Illustration of TAD clusters identified using proliferating (*left*) and senescent HUVEC Hi-C data (*right*) for chr10 and 14. Spheres represent TADs; the most 5'/3' TADs (*purple* and *yellow dots*, respectively) and "singular" TADs (*arrows*) are indicated.

(C) Line plots showing average HMGB1 (*green*), CTCF (*grey*), and H3K27ac ChIP-seq profiles (*magenta*) at the extremities of all clustered TADs (±20 kbp; *top*), of clusters of >3 TADs (*middle*) or of "singular" TADs (*bottom*) from proliferating IMR90.

(D) Exemplary Hi-C data (40-kbp resolution) around a "singular" TAD (*magenta*) in HUVEC chr10 aligned to TAD positions, HMGB1 peaks, and ENCODE ChIP-seq.

(E) As in panel D, but for TADs that form a >3-TAD cluster in TiLO data.

(F) Heatmap showing senescence-induced changes in expression levels ($log_2$FC) of SASP-related genes embedded in singular TADs like those in panel B. Genes bound by HMGB1 are indicated (arrows), and are not morethan what would be expected by chance; $P$>0.1, Chi-square Goodness-of-Fit test.

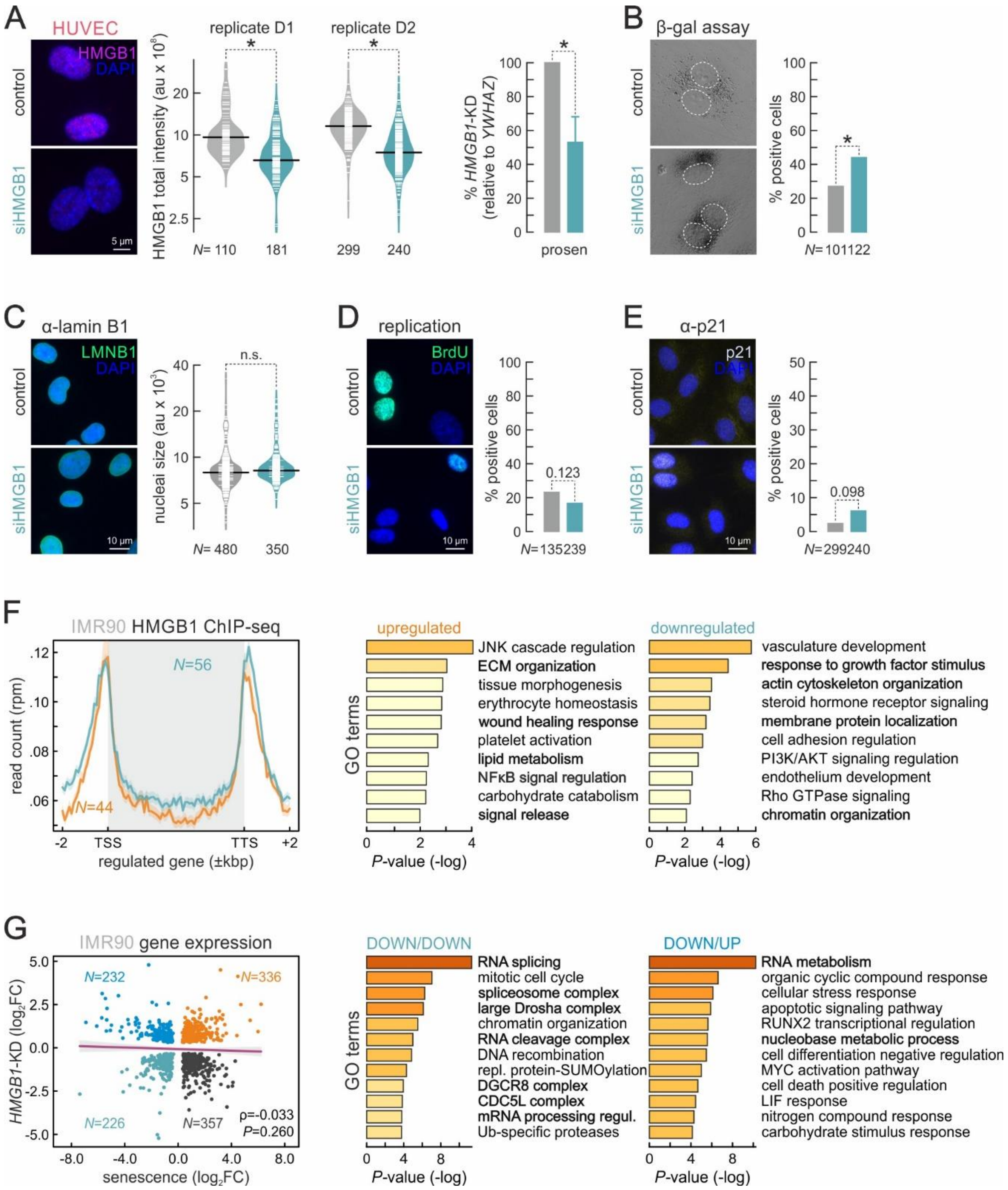


**Figure S6. Effects of *HMGB1* knockdown in proliferating HUVECs and IMR90.**

(A) Representative immunofluorescence images of siHMGB1-treated HUVECs showing reduced HMGB1 levels compared to control cells (*left*). Bean plots quantify knockdown efficiency (*middle*; *N* indicates the number of cells analyzed). Bar graphs (*right*) show normalized *HMGB1* mRNA levels (± S.D.; *N*=2) in knockdown compared to control cells. Bar: 5 µm. *: *P*<0.01; Wilcoxon-Mann-Whitney and unpaired two-tailed Student's t-test for bean and bar plots, respectively.

(B) Representative brightfield images of siHMGB1-treated HUVECs showing elevated β-galactosidase activity compared to control cells (*left*); bar graphs quantify this increase (*right*; *N* indicates the number of cells analyzed). *: *P*<0.01; Fisher's exact test.

(C) As in panel A, but for LMNB1 levels. Bar: 10 µm. No statistically significant difference (n.s.); Wilcoxon-Mann-Whitney test.

(D) As in panel A, but for BrdU incorporation. Bar: 10 µm. *P*=0.123; Fisher's exact test.

(E) As in panel A, but for p21 levels. Bar: 10 µm. *P*=0.098; Fisher's exact test.

(F) Line plots (*left*) showing average HMGB1 ChIP-seq profiles along genes up- (*orange*) or downregulated upon *HMGB1*-knockdown in IMR90 (*green*; *N* indicates the number of directly HMGB1-bound genes). Bar plots (*middle*/*right*) showing GO terms associated with either gene set and their enrichment *P*-values; terms relevant to senescence are highlighted (*black*).

(G) Scatter plots (*left*) showing correlation between differentially-expressed genes ($log_2$FC) upon senescence entry and *HMGB1*-knockdown in IMR90. The Pearson's correlation (ρ) and its associated *P*-value are shown alongside the number of genes (*N*) in each subgroup. Bar plots (*middle*/*right*) showing GO terms and enrichment *P*-values associated with genes commonly downregulated ("DOWN/DOWN") or downregulated in senescence but upregulated upon *HMGB1*-knockdown ("DOWN/UP"); terms relevant to RNA processing are highlighted (*black*).

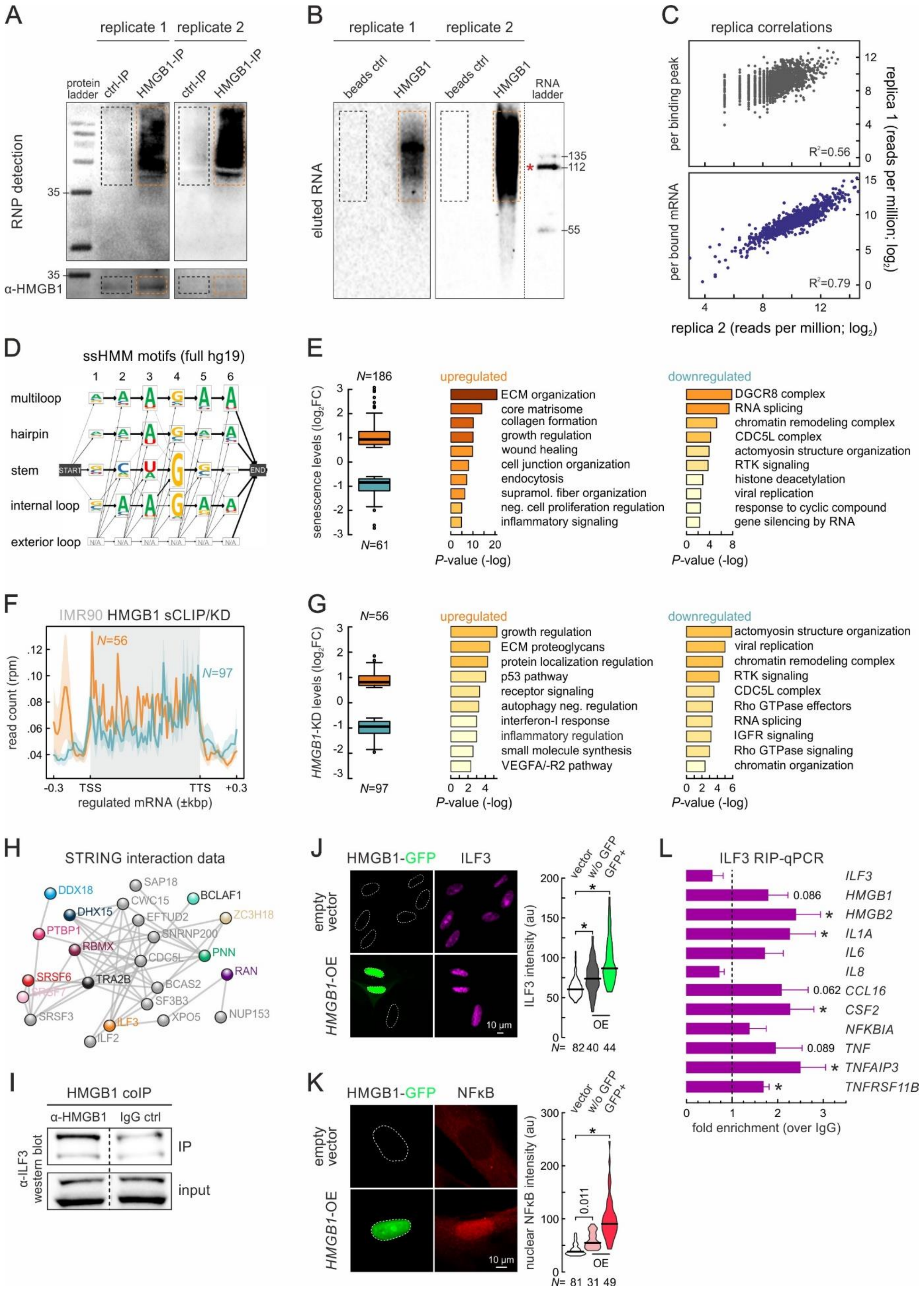
A
replicate 1
replicate 2
protein ladder
ctrl-IP
HMGB1-IP
RNP detection
α-HMGB1
B
beads ctrl
HMGB1
RNA ladder
eluted RNA
C
replica correlations
per binding peak
per bound mRNA
R²=0.56
R²=0.79
replica 1 (reads per million; log2)
replica 2 (reads per million; log2)
D
ssHMM motifs (full hg19)
multiloop
hairpin
stem
internal loop
exterior loop
E
senescence levels (log2FC)
N=186
N=61
upregulated
ECM organization
core matrisome
collagen formation
growth regulation
wound healing
cell junction organization
endocytosis
supramol. fiber organization
neg. cell proliferation regulation
inflammatory signaling
downregulated
DGCR8 complex
RNA splicing
chromatin remodeling complex
CDC5L complex
actomyosin structure organization
RTK signaling
histone deacetylation
viral replication
response to cyclic compound
gene silencing by RNA
P-value (-log)
F
IMR90 HMGB1 sCLIP/KD
N=56
N=97
read count (rpm)
regulated mRNA (±kbp)
G
HMGB1-KD levels (log2FC)
upregulated
growth regulation
ECM proteoglycans
protein localization regulation
p53 pathway
receptor signaling
autophagy neg. regulation
interferon-I response
inflammatory regulation
small molecule synthesis
VEGFA/-R2 pathway
downregulated
actomyosin structure organization
viral replication
chromatin remodeling complex
RTK signaling
CDC5L complex
Rho GTPase effectors
RNA splicing
IGFR signaling
Rho GTPase signaling
chromatin organization
H
STRING interaction data
I
HMGB1 coIP
α-HMGB1
IgG ctrl
IP
input
α-ILF3 western blot
J
HMGB1-GFP
ILF3
empty vector
HMGB1-OE
vector
w/o GFP
GFP+
ILF3 intensity (au)
OE
K
HMGB1-GFP
NFκB
nuclear NFκB intensity (au)
L
ILF3 RIP-qPCR
ILF3
HMGB1
HMGB2
IL1A
IL6
IL8
CCL16
CSF2
NFKBIA
TNF
TNFAIP3
TNFRSF11B
fold enrichment (over IgG)

Figure S7. HMGB1 sCLIP controls and analysis of HMGB1-interacting RBPs.

(A) Electrophoretic profiles of control (beads only; *black dotted square*) and HMGB1 IP (*orange dotted rectangle*) probed for RNA (*top*) or the HMGB1 protein (*bottom*) in both sCLIP replicates. The 35-kDa band of the molecular mass ladder is indicated.

(B) Electrophoretic profiles of RNA eluted from control (beads only; *black dotted square*) or HMGB1 IP (*orange dotted rectangle*) in both sCLIP replicates. The 112-nt band of the molecular mass ladder is indicated (*red star*) and corresponds to ~144 pg of RNA.

(C) Scatter plots showing correlation of sCLIP data from two independent biological replicates compared per binding peak (*top*) or per bound mRNA normalized read count (*bottom*). Spearman correlation values ($R^2$) are indicated.

(D) Output of ssHMM motif analysis of sCLIP data showing sequence probabilities in HMGB1-bound motifs predicted to form different structures using the reference human genome (hg19).

(E) Box plots (*left*) showing expression fold-changes ($log_2$) of mRNAs differentially-regulated upon senescence and bound by HMGB1 in IMR90 sCLIP. The number of peaks (*N*) analyzed is indicated below each bar. Bar graphs (*right*) with GO terms associated with each subgroup and their enrichment *P*-values are also shown.

(F) Line plots (*left*) showing average HMGB1 sCLIP profiles along mRNAs up- (*orange*) or downregulated upon *HMGB1*-knockdown in IMR90 (*green*; *N* indicates the number of directly HMGB1-bound genes).

(G) As in panel E, but for the up-/downregulated genes from panel F.

(H) Experimentally-validated protein-protein interaction network for HMGB1 co-immunoprecipitating RBPs (*colored spheres*) that are also downregulated in senescence. Their secondary interactors (*grey spheres*) from the STRING database are also shown.

(I) Western blots using antisera recognizing ILF3 on proteins co-immunoprecipitating with HMGB1 (*left*) or IgG (*right*) in IMR90; blots on 10% of input material provides a loading control.

(J) Representative images of IMR90 overexpressing HMGB1-GFP and immunostained for ILF3. Nuclear outlines are indicated (*dashed lines* based on DAPI counterstaining). IMR90 transfected with empty vectors provide a control. Bar: 10 μm. Signal intensities are quantified in the violin plots (*right*). *: significantly different to control; $P<0.001$, Wilcoxon-Mann-Whitney test

(K) As in panel I, but immunostained for phospho-NFκB. Bar: 10 μm.

(L) Bar graphs showing mean fold-enrichments (over IgG controls ±S.D., $N=2$) of selected mRNAs from ILF3 RIP experiments in proliferating IMR90. *: significantly different to control; $P<0.05$, unpaired two-tailed Student's t-test

Supplemental Table Legends (tables are provided as .xlsx files online)

**Table S1. HMGB1 sCLIP target mRNAs.** List of high-confidence HMGB1-bound mRNAs from two sCLIP biological replicates.

**Table S2. HMGB1 ChIP-seq peaks and long-range loops.** List of high confidence HMGB1 ChIP-seq peaks from (A) proliferating HUVEC and (B) proliferating IMR90. (C) List of intrachromosomal HMGB1 long-range loops from proliferating HUVEC.

**Table S3. Differentially-expressed genes.** Differentially-expressed genes between (A) all clusters in single cell nascent RNA-seq data; (B) proliferating and senescent HUVEC; (C) proliferating and senescent IMR90; (D) control and *HMGB1*-knockdown IMR90.

**Table S4. HMGB1 interacting proteins.** Full list of proteins (A) co-immunoprecipitating with HMGB1 (significant ones highlighted in bold), alongside a summary of the biological replicates submitted, and the technical parameters of the proteomics run; (B) recovered by whole-cell proteomics comparing proliferating to senescent IMR90.

**Table S5. Ribo-seq data analysis.** Full list of genes tested in Ribo-seq (performed in biological triplicates) alongside their total RNA, translation, and buffering indexes.
